# Molecular mechanism of Na^+^/H^+^ antiporting in NhaA

**DOI:** 10.1101/2024.11.23.624977

**Authors:** Tengyu Xie, Jiahao He, Shuo Sun, Yulei Chen, Jing Huang, Yandong Huang

## Abstract

Sodium-proton antiporter NhaA of *Escherichia coli* is a paradigm to investigate the mechanistic basis of the fundamental Na^+^/H^+^ exchange in cells. However, all existing crystal structures of NhaA are inward-facing (IF) and the putative outward-facing (OF) structures are still unsolved by experiment, limiting a complete understanding of the transport cycle where Lys300 plays a key role in both structural stability and transport function. Here, we report a set of regular molecular dynamics (MD) simulations that start from the structure predicted by an artificial intelligence method that generates function-relevant alternative conformations. It is found that NhaA rapidly relaxes into either the IF or OF conformation. Furthermore, neutralization of Lys300 allows two sodium ions bound to the reaction cavity, which is associated with enhanced conformational sampling. Based on these observations, we propose a sodium-coupled mechanism of Na^+^/H^+^ antiporting.

---

The exchange of sodium ions and protons across cellular membranes by Na^+^/H^+^ antiporters is essential for the homeostasis of pH, Na^+^ concentration and cell volume ^1, 2^. Na^+^/H^+^ antiporters are well-established drug targets of a variety of human diseases, including hypertension ^3^, diabetes ^4^ and cancer ^5–7^, and indispensable for salt resistance in plants, such as Pyropia haitanensis^8^. Among these antiporters, NhaA of *Escherichia coli* has been studied extensively as a prototype both experimentally and theoretically ^9^. NhaA functions by exchanging two protons in the periplasm for one sodium in the cytoplasm, exhibiting an electrogenic transport ^10, 11^. Na^+^/H^+^ antiporting is characterized by dramatic pH dependence. For instance, NhaA is inactive below pH 6.5 and the turnover rate increases by over 3 orders of magnitude from pH 6.5 until pH 8.5 at which the maximal is reached ^12^. NhaA is a homodimer in nature ^13^ but monomers are fully functional ^14^. From the crystal structure solved at inactive pH 4.0, each monomer contains a stationary dimer domain that connects two monomers and a dynamic core domain that contains a cavity in the middle of the membrane for substrate ion binding (Fig. 1A) ^15^. It has been demonstrated that the conformational dynamics of NhaA increases with pH, matching the pH-dependent transport activity ^16–21^.

**Figure 1:**
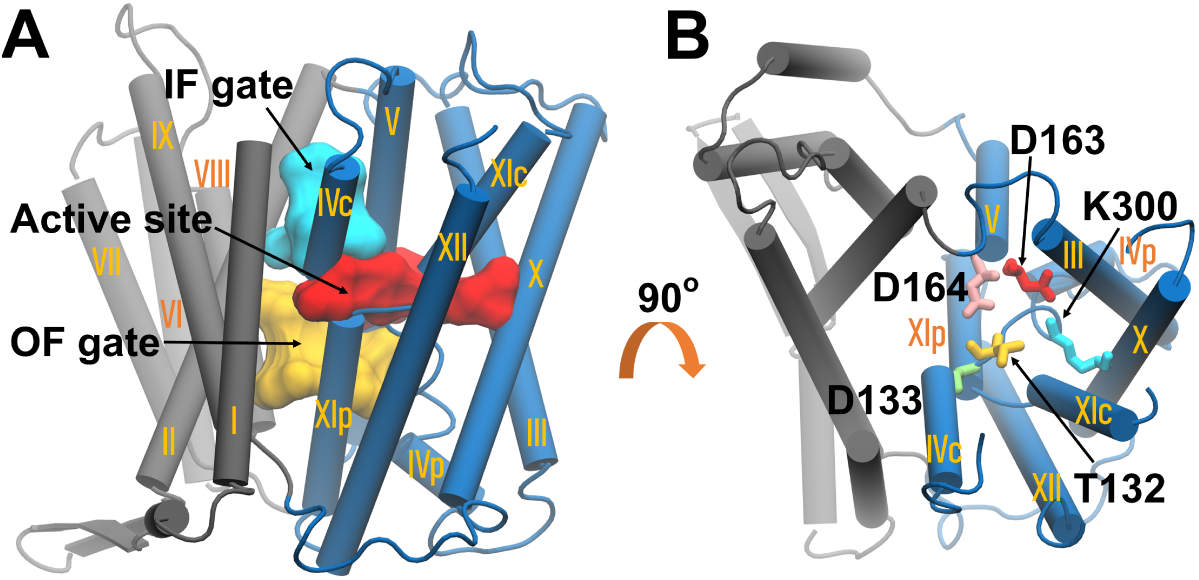
Crystal structure of NhaA (PDB code: 4AU5). Secondary structures are displayed with the cartoon model. A Sideview of NhaA with the cytoplasmic side at the top. The 12 transmembrane helices (TMs) are indexed with roman numerals, namely 12–30 (I), 59–85 (II), 95–116 (III), 121–131 (IVp), 134–143 (IVc), 150–175 (V), 182–200 (VI), 205–218 (VII), 223–236 (VIII), 247–271 (IX), 290–311 (X), 327–336 (XIc), 340–350 (XIp) and 357–382 (XII) 15. Core and dimer domains are colored blue and gray, respectively. The IF and OF gates are presented by the surface model and colored cyan and orange, respectively. IF gate is composed by V75, I134, M157, A160 and I161. OF gate includes F72, A167, I168, F344 and I345. The active site that locates in the middle of IF and OF gates is colored red and composed by T132, D133, D163, D164 and K300. **B** Top view of NhaA. Active site residues are shown with the stick model and colored orange (T132), green (D133), red (D163), pink (D164), and cyan (K300), respectively.

To understand the transport mechanism, it is of importance to identify the sodium binding sites as well as proton carriers within the reaction cavity of the core domain. Thr132, Asp133, Asp163, Asp164 and Lys300 (Fig. 1B) that are evolutionary conserved and locate in the cavity have been suggested to form the active site of NhaA ^22^. ITC experiments showed that Asp163, Asp164, and the backbone of Thr132 and Asp133 are substantial parts of the NhaA cation binding site, though the observed ITC signals were obtained using Li^+^, an alternative cation transported by NhaA ^23^. Later it was suggested that a sodium ion is mostly bound to the backbone of Thr132 and side-chain carboxyl oxygens of Asp163 and Asp164, excluding Asp133 from the putative sodium binding sites ^24^. More recently, the active site was found capable of accomodating two sodium ions when Lys300 is deprotonated or neutralized ^25^. However, it is unclear such a twin sodium binding has any physiological relevance.

It has been a consensus that Asp164 is one of the two proton carriers ^24, 26^. However, the second proton carrier is still a controversy. Asp163 was proposed as the second proton since the first high-resolution crystal structure (PDB code: 1ZCD) measured at inactive pH 4.0 ^26^. A MD study proposed that Asp163 acts as a conformational switch by changing its protonation state ^27^. The idea was later challenged by the second crystal structure (PDB code: 4AU5) measured at pH 3.8 where Asp163 forms a stable salt bridge with Lys300 (Fig. 1B), a feature absent in the earlier structure ^28^. Recently, the presence of the salt bridge was further validated by the third crystal structure (PDB code: 7S24) solved under active pH 6.5, although with only 10% of the wide type activity ^25^. Because of the salt bridge, Lys300 and Asp163 are thought to be positively and negatively charged, respectively. Consequently, Lys300, instead of Asp163, was identified as the second proton carrier ^28^. It was suggested that Lys300 releases the proton upon sodium binding that interrupts the salt bridge, which leads to a new mechanism that highlights the competition between sodium and proton in the active site ^24^.

The debate over whether Lys300 is the second carrier continues. NapA from *Thermus thermophilus* is a homology of NhaA. Mutation of Lys305, the equivalent residue of Lys300 in NhaA, to arginine, switches NapA from being electrogenic to electroneutral, supporting Lys300 of NhaA as a proton carrier ^29^. In addition, the mammalian Na^+^/H^+^ exchanger NHA2, also homologous to NhaA, contains two aspartate residues Asp277 and Asp278, which are counterparts of Asp163 and Asp164 in NhaA. It was found that NHA2 performs electroneutral rather than electrogenic transport, suggesting that the proton is carried the most by one aspartate, most likely the Asp278, which from a mechanistic point of view supports Lys300 of NhaA as the second proton carrier ^30^. Further, a phylogenetic analysis identified a minimum of four electrogenic clades, including NhaA. The phylogeny suggested that these electrogeneic transporters are characterized by an aspartate and a lysine in the active site ^11^, which echoes with above two electrophysiological studies. More straightforwardly, replacing Lys300 of NhaA with other residues, low thermal stability and reduced activity were detected ^31^. Besides, Lys300 was shown to be important in pH regulation ^23, 32, 33^. Nevertheless, the electrogentic transport was maintained when Lys300 was mutated to an uncharged residue, like alanine ^31^ and glutamine ^34^, which favors Asp163 as the second proton carrier. Interestingly, mutating Asp163 and Lys300 to asparagine and glutamine, respectively, still produced an electrogenic variant ^34^, implying that neither Asp163 nor Lys300 is essential for electrogentic transport of NhaA. To reconcile the contradiction, a third mechanism was proposed where Asp163 takes the role of Lys300 as the second proton carrier once Lys300 is mutated ^31^. Such a back-up proton carrier hypothesis was updated later by a MD study where Lys300 mutants offer the electrogenic transport by using an alternative proton-binding residue Asp133 ^35^.

Release of the first proton by Asp164 has been suggested to trigger IF gate (Fig. 1A) opening and the resulting sodium bound to the active site ^24^. In fact, NhaA adopts the conceptual alternating access model ^2^, which asks for an OF gate (Fig. 1A). Meanwhile, it is of importance to see whether the second proton released presumably by Lys300 is coupled with the subsequent transition from IF to the putative OF conformation. However, existing crystal structures of NhaA are all IF ^15, 25, 28^, thus accurate prediction of OF conformation is crucial. To switch between two states, NhaA has to overcome a barrier, which is 4.8 kcal/mol probed by experiment ^9^ and 14 to 16 kcal/mol estimated by theory ^36^. Apparently, such a conformational change is hardly driven by the thermal energy, about 0.6 kcal/mol at 300 Kelvin, during a regular MD simulation, asking for enhanced sampling.

Based on transition-path shooting simulations, the OF conformation of a homology of NhaA, namely PaNhaP from archaea *Pyrococcus abyssi*, was obtained ^40^. However, during the allosteric process, which motion is essential was not addressed. On the other hand, an OF structure of NhaA was predicted by metadynamics simulations where conformational sampling was biased along an angular motion ^41^. It was found that NhaA employs roughly the same rocking-bundle and elevator mechanism with NapA where the initial angular movement is sufficient to induce the alternate access ^41^. However, the essential sodium-proton exchange was seldom incorporated by enhanced sampling schemes above. Earlier, Arkin et al proposed a mechanism of Na+/H+ antiporting by simulating NhaA under four possible combinations of the protonation states of Asp163 and Asp164 in the presence of sodium ions ^27^. Though a conformational change in response to the protonation of Asp163 was detected, the global conformation was not affected, in contradiction to the elevator-like movement which has been validated extensively by experiment ^37–39^ and theory ^40, 41^. Likewise, four MD simulations with different protonation states selected from the titration space of Asp163, Asp164 and Lys300 were set out to discuss proton-coupled sodium binding in the framework of IF state, which nevertheless lacks any characterizations under the OF state ^25, 28^.

To unlock the OF conformation of NhaA, forty simulations were initialized from one of the potentially alternative structures generated by the multi-conformation prediction method AFEXplorer (AFEX) ^42^. Further, to investigate possible effects of neutralizing Lys300 on the structural stability and function of NhaA, simulations were divided into two groups that correspond to deprotonated and protonated Lys300, respectively. The study shows that simulations can relax towards between IF and OF states. More importantly, deprotonation of Lys300 allows the binding of two sodium ions in the active site, which would accelerate the transition kinetics between IF and OF conformations. Together, our work shows how changes in protonation state of Lys300 contribute to sodium binding and state transition.

## Results

### Overview of simulations

We began with the IF crystal structure (PDB code: 4AU5) to initialize two ten-microsecond-timescale regular simulations, where the OF state was not sampled. To uncover the OF conformation currently inaccessible by experiment, we used AFEX to explore potential alternative states and selected one to initialize forty independent runs, each of which lasted 500 nanoseconds, except for the four extended to 2 microseconds. Notably, to study effects of neutralizing Lys300, above two kinds of simulations were divided into two groups that correspond to protonated and deprotonated states of Lys300, respectively. In addition, considering Asp164 as a proton carrier, eight simulations that lasted 2 microseconds each were carried out, in which both Asp164 and Lys300 were protonated. Convergence analyses are given in Supplementary Fig. 1-5. We note that analyses of simulation trajectories below are based on the forty independent runs, except for those that concern sodium binding in IF.

### OF gate opens

Features of the AFEX-predicted structure (Supplementary Structure 1) was inspected by aligning its dimer to that of the crystal structure (PDB code: 4AU5). Fig. 2A shows that the cytoplasmic side of core is rotated by about 20 degrees, which results in transmembrane helices (TMs) IVc and XII of core approaching to TMs I and II of dimer, respectively. As illustrated in Fig. 2B, the OF gate residues, especially Ile168 and Ile345 in core, are affected by such a rotational motion. For instance, the minimal distance between Ile168 and Phe72 of dimer increases from 3.7 to 5.2 Å . Besides, Ile345 is displaced away from F72 by 1.4 Å . Finally, Ile168 and Ile345 are separated by 4.7 Å (Supplementary Table 1). Apparently hydrophobic interactions in OF gate have been weakened in the predicted structure, which might accelerate OF gate opening.

**Figure 2:**
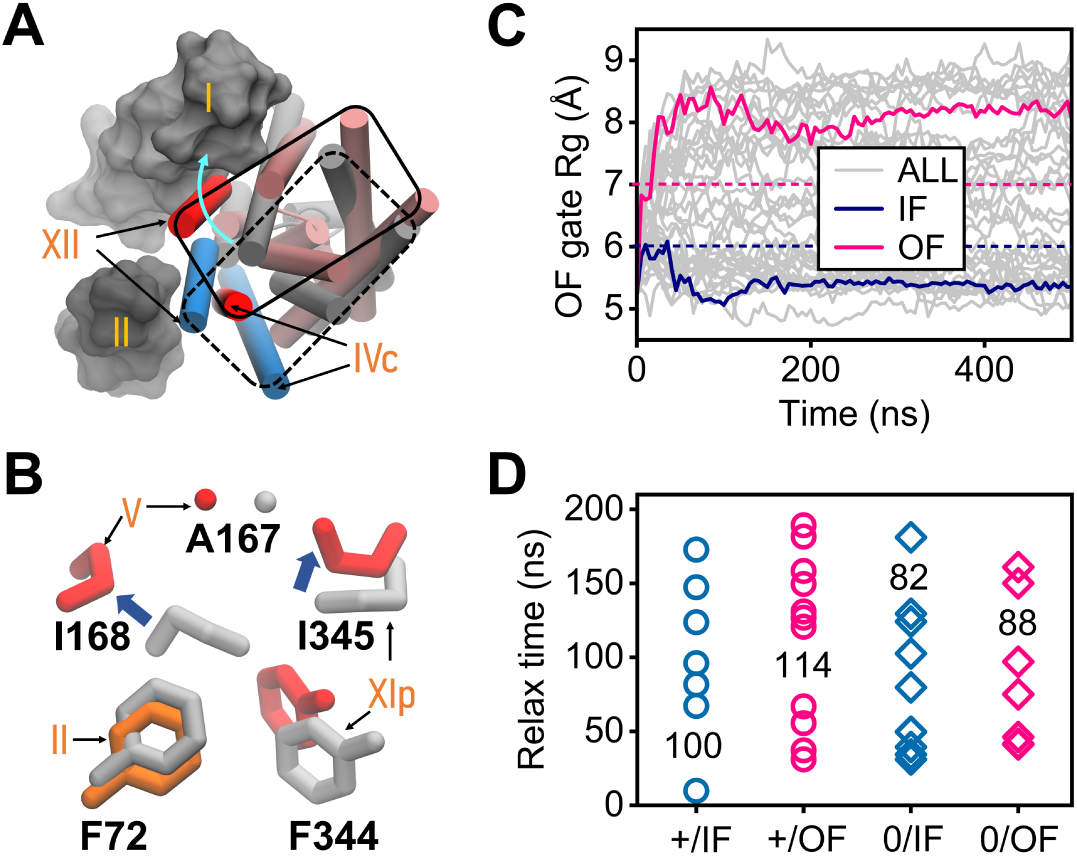
Simulations initialized from the AFEX-predicted structure. **A** Dimer of the predicted structure aligned to that of the crystal one (PDB code: 4AU5). TMs I and II that belong to the dimer are displayed with the surface model. Cores (TMs IVc and XII of the core) of the predicted and crystal structures are displayed with the cartoon model and colored pink and gray (red and blue), respectively. Solid (predicted) and dashed (crystal) frames represent the border of core at cytoplasm. The cyan arrow indicates the rotation from dashed to solid frame. **B** Side chains of OF gate residues F72 on TM II, A167 and I168 on TM V and F344 and I345 on TM XIp, for the crystal (gray) and predicted (red except for F72 in orange) structures, respectively. The blue arrows indicate the displacements of I168 and I345 with respect to F72. **C** Radius of gyration (Rg) of OF gate as a function of time (ns). OF gate Rg above 7.0 Å (dashed red line) or below 6.0 Å (dashed blue line) is defined as OF or IF state. The background color gray is applied to all trajectories, among which two runs that converged to OF and IF are highlighted with solid red and blue lines, respectively. **D** OF gate Rg relax time are divided into four groups, K300 charged in IF (+/IF) and OF (+/OF) and neutralized in IF (0/IF) and OF (0/OF). The average for each group is displayed.

Indeed, OF gate opening was observed in simulations that started from the AFEX-predicted structure (Fig. 2C). Encouragingly, both backward IF (Fig. 2C) and forward OF states can be achieved, implying that AFEX might help overcome the barrier between OF and IF states. Here cutoffs 6 and 7 Å of OF gate Rg were utilized to distinguish OF from IF state. In specific, IF state corresponds to OF gate Rg below 6 Å whereas OF state above 7 Å . To understand whether the equilibration is coupled with the protonation state of Lys300, auto-correlation of OF gate Rg was calculated (Fig. 2D). The gate is considered relaxed once auto-correlation reaches zero. Only simulations that converged to IF or OF were considered. As a result, 7 and 11 simulations converged to IF and OF, respectively, when Lys300 is charged. In contrast, 10 and 7 runs relaxed to IF and OF, respectively, where Lys300 is in neutral state. Obviously, it took the OF gate less than 200 ns relaxing to either IF or OF state. Notably, the mean relax time reduces from 100(114) to 82(88) ns for IF(OF) when Lys300 is neutralized.

### Deprotonation of Lys300 enhances conformational dynamics

It’s noticed from Fig. 2C that forty independent simulations provide a scattered sampling of OF gate Rg above 7 Å, which implies a rugged energy profile in OF state. As a result, free energy maps of IF versus OF gate Rg were plotted. Here the probability density was converted to relative free energy ΔG calculated by -RTln(n*_ij_*/N), where R is the Boltzmann constant, T denotes room temperature of 300 Kelvin, n*_ij_* is the number of data points in the bin, and N is the total number of data points collected in the entire region. As illustrated in Fig. 3A where Lys300 is charged, there are three distributions in OF. It is worth noting that when OF gate is wide open (OF gate Rg *>* 8 Å), conformations with IF gate Rg around 4.5 (OF-a) and 5.5 (OF-b) Å were sampled. OF-a is more populated than OF-b when Lys300 is charged. Whereas a single minimal is obtained at IF side when OF gate is closed (OF gate Rg *<* 6 Å). Compared to the crystal structure, the increment of either IF or OF gate Rg (∼0.5 Å) in the AFEX-predicted structure is observed.

**Figure 3:**
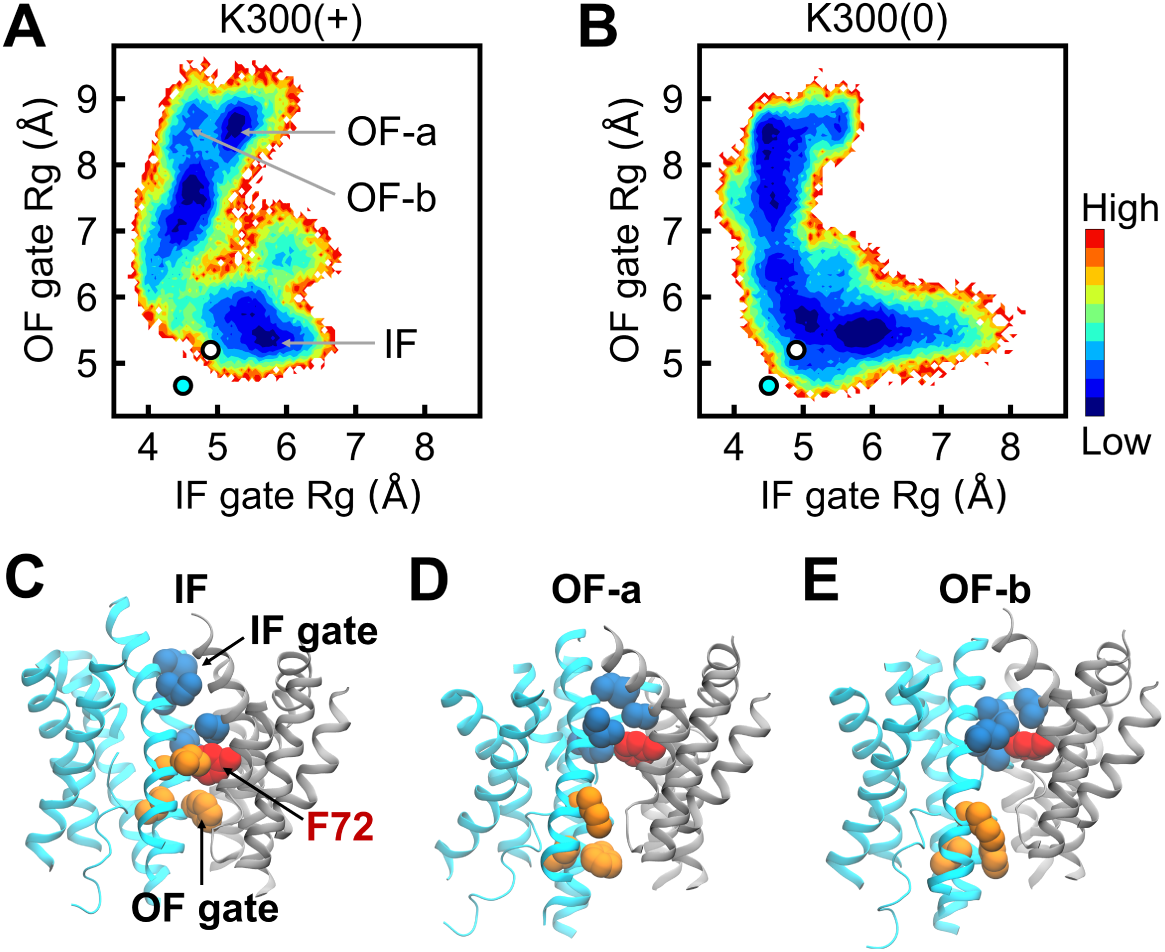
Effects of neutralizing Lys300 on gating dynamics. **A** Free energy maps of IF and OF gate Rg for charged K300 or K300(+). The energy minimal below 6 Å of OF gate Rg is defined as the IF state. Two energy minimals in the range from 8 to 9 Å of OF gate Rg are defined as OF-a and OF-b, respectively. AFEX-predicted and crystal (PDB code: 4AU5) structures are marked with white and cyan circles, respectively. **B** Free energy maps of IF and OF gate Rg for K300 neutralized or K300(0). **C** Side view of a snapshot that shows open IF gate and closed OF gate in IF state. Dimer and core are displayed with the new cartoon model and colored gray and cyan, respectively. Side chains of IF and OF gates are displayed with spheres and colored blue and orange, respectively, expect for red-colored F72 of OF gate. **D** A snapshot with open OF gate and half open IF gate in OF-a. **E** A snapshot with open OF and closed IF gates, respectively, in OF-b.

Interestingly, as illustrated in Fig. 3B, the energy surface is flattened upon the deprotonation of Lys300, especially the area in the middle of IF and OF which is hardly accessible in Fig. 2A. Such smoothed energy barriers would accelerate the sampling in conformation. Besides, populations of OF-a and OF-b are reversed and OF-b even prevails or is the most stable conformation in OF state, exhibiting strict alternating gate opening. At IF side where OF gate Rg is below 6 Å, IF gate is more open when compared with that in Fig. 3A, in consistence with constant pH MD (CpHMD) simulations ^24^ and the present simulations initialized from the crystal structure (Supplementary Fig. 6).

Looking into a snapshot structure of IF in Fig. 3C, IF and OF gates are open and closed, respectively. OF gate residue Phe72 is positioned in the middle of IF and OF gates. From panels D and E of Fig. 3, OF gate opening is characterized by the displacement of other OF gate residues away from Phe72 and towards periplasm. On the other hand, IF gate can be half (OF-a, Supplementary Structure 2) or fully (OF-b, Supplementary Structure 3) closed. In contrast to other OF gate residues with evident conformational changes, Phe72 that belongs to dimer is unchanged. Thus, Phe72 may act as a barrier that controls the motion of core with respect to dimer.

### Validation of the simulated OF state

It has been suggested that a rotational motion of core with respect to dimer is essential for the conformational change from IF to OF ^41^. In this work, ΔΨ, an angle between the two domains, is defined to probe the rotational motion (Fig. 4A) ^41^. Encouragingly, ΔΨ of the putative OF conformations in our simulations is larger than that of the crystal or AFEX-predicted structure by about 20 degrees (Fig. 4B), which is consistent with the OF states obtained from metadynamics simulations ^41^. The angular motion is accompanied by an elevator-like movement which can be measured by ΔZ (Fig. 4C) that represents the displacement of core with respect to dimer along Z axis ^41^. As illustrated in Fig. 4D, core moves towards periplasm by about 5 Å from IF to OF, which is also in line with metadynamics simulations. We note that the distribution at IF side for either ΔΨ or ΔZ resembles that by simulations starting from the crystal structure (Supplementary Fig. 7).

**Figure 4:**
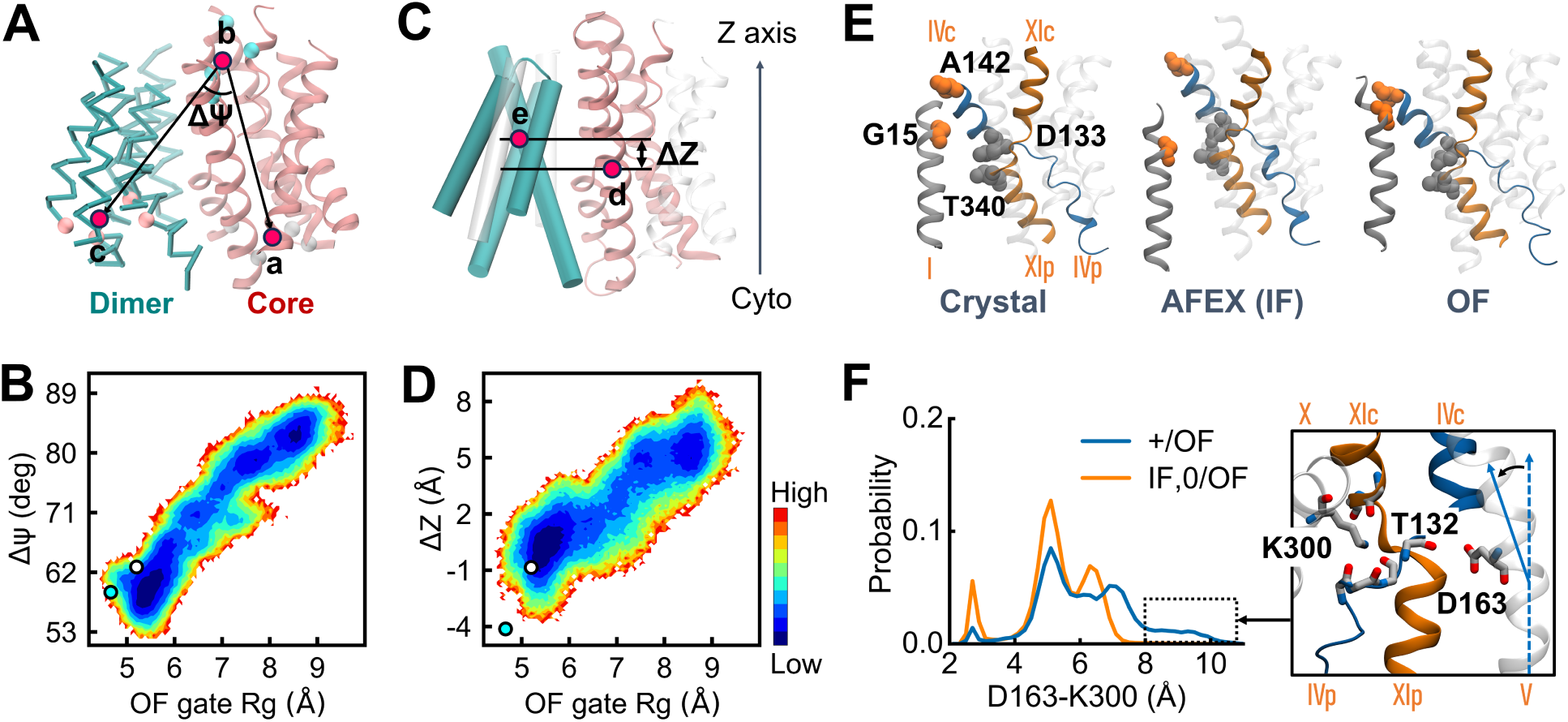
Conformational change in NhaA. **A** Definition of the angle ΔΨ between core and dimer by three end points a, b and c, each of which is the center of mass of four C*_α_* atoms. a is defined using residues E124, I171, A346 and L357, b by L135, S146, T323 and L371 and c by L77, T205, G234 and L251. Presented is the side view of NhaA where the bottom is the cytoplasmic side (Cyto). Dimer (green) and core (pink) domains are displayed with the trace and new cartoon models, respectively. **B** Free energy map as a function of OF gate Rg and ΔΨ. AFEX-predicted and crystal structures are marked with white and cyan circles, respectively. **C** Definition of the Z-axis distance ΔZ between core and dimer domains by d and e, z coordinates of centers of mass of two four-helix bundles. d is defined by residues T122 to R147, V148 to Y175, Q326 to V353 and D354 to R383 (pink) and e by residues L60 to Q85, T205 to K221, S222 to F236 and P247 to A272 (green). The rest helices are transparent. ΔZ equals d minus e. Dimer and core domains are represented by the cartoon and new cartoon models, respectively. **D** Free energy map as a function of OF gate Rg and ΔZ. **E** Crystal structure (PDB code: 4AU5) is displayed with the new cartoon model. TMs I, IV and XI are colored gray, orange and blue, respectively. G15, A142 (orange), D133 and T340 (gray) side chains shown with spheres are apart from each other in 4AU5. D133 contacts with T340 in the AFEX-predicted structure (AFEX) or IF from simulations. In addition to the contact above, A142 is in contact with G15 in OF. **F** Probability distributions of the minimal distance between D163 and K300 side chains (D163-K300) for charged K300 in OF state (+/OF) (blue) and the rest (orange), including IF and K300 neutralized in OF (0/OF). A snapshot showing, in the region of 8 Å and above, the interaction between K300 and the cleft created by TMs IVp and XIc, bending of TM V and dissociation of D163 from T132 backbone. D163 on bent TM V, K300 on TM X and backbones of P129, A130, A131 and T132 on TM IVp (blue) and G332 and C335 on TM XIc (orange) are displayed with sticks where oxygens, nitrogens and carbons are colored red, blue and gray, respectively. The dashed and solid lines indicate straight and bent TM V, respectively. Helices are displayed with the new cartoon model.

Two initial structures are marked in Fig. 4B and Fig. 4D so as to examine their difference, if any, in ΔΨ or ΔZ. It is noticed that ΔΨ of the AFEX-predicted structure (Fig. 4B) is closer to OF but not trapped in the minimal of IF, making it easier to evolve to OF. On the other hand, ΔZ for the predicted structure (Fig. 4D) is more likely stuck in IF. Thus, the increase of ΔΨ made by AFEX, though small, could be more important than that of ΔZ, matching the conclusion of metadynamics simulations ^41^. From paths of ΔZ and ΔΨ (Supplementary Fig. 8), it is found they move together from IF to OF rapidly when Lys300 is charged. Whereas, once Lys300 is neutralized, they stay temporarily in an intermediate state before reaching OF.

Apart from above global conformational changes, contacts between Gly15 and Ala142 ^41^ and between Asp133 and Thr340 ^43^ under OF have been probed by experiment. As illustrated in Fig. 4E, both contacts are broken in the crystal structure (PDB code: 4AU5). The contact between Asp133 and Thr340 is present in the AFEX-predicted structure and dominant in the simulated IF. (Supplementary Fig. 9). Encouragingly, both contacts were reproduced in the current simulated OF (Supplementary Fig. 10), indicating its reasonability.

While the salt bridge between Asp163 and Lys300 forms in the two recent crystal structures (PDB codes: 4AU5 ^28^ and 7S24 ^25^) and breaks in the first one (PDB code: 1ZCD), IF-based investigation has covered its role in structural stability or sodium-proton exchange ^24^. However, it is still unknown in OF. To explore possible influence of this hallmark interaction on IF-OF transition, the minimal distance between Asp163 and Lys300 side chains was calculated. As shown in Fig. 4F, the distribution for charged Lys300 in OF (+/OF) is diffusive when compared with others. Particularly only +/OF offers the distance larger than 8 Å . Interestingly, the bending of TM V as well as the dissociation of Asp163 sidechain from T132 backbone (Supplementary Structure 2) was observed (Fig. 4F) when the distance between Asp163 and Lys300 is above 8Å (Supplementary Fig. 11), consistent with previous CpHMD and MD simulations in IF where Lys300 is deprotonated ^24^. Instead of the intact salt bridge that prevails in the simulation initialized from the crystal structure where Lys300 is charged (Supplementary Fig. 12), the bulk population in Fig. 4F, regardless of the protonation state of Lys300, ranges from 4 to 8 Å, which nevertheless agrees with CpHMD simulations at alkaline pH side ^24^. It is worthwhile noting that the minimal distance between Lys300 and Asp163 was increased from 2.5 to 3.6 Å at membrane normal by AFEX (Supplementary Fig. 13). Thus, the low ratio of salt bridging between Lys300 and Asp163 could be explained by the fact that Lys300 side chain, as illustrated in Fig. 4F, is stabilized via inserting into a negatively charged cleft formed by helical terminals of TMs IVp and XIc (Supplementary Fig. 14).

### IF and OF funnels are solvated alternately

Water molecules are supposed to enter the active site when a funnel is open ^24^, which is indispensable for the sodium-proton exchange via IF and OF funnels. As illustrated in Fig. 5A, the active site that contains four ionizable residues locates nearby the middle of lipid bilayer at z coordinate of zero. Water number across the membrane was computed (Fig. 5B), from which one can see that IF funnel opens and simultaneously OF funnel is blocked in IF state and vice versus. The active site around z coordinate of zero is accessible by solvent alternately, which can be further visualized using water density profiles of IF (Fig. 5C) and OF (Fig. 5D), respectively. Notably, as illustrated in Fig. 5E, a continuous water chain, though unstable, can be found at the early stage t_1_ of the transition to OF at t_2_ (Supplementary Movie 1). In particular, water molecules from both sides are accessible to Asp163 or Asp164 at t_1_. Whereas, water molecules from cytoplasm are excluded at t2, consistent with above water number and density calculations for OF.

**Figure 5:**
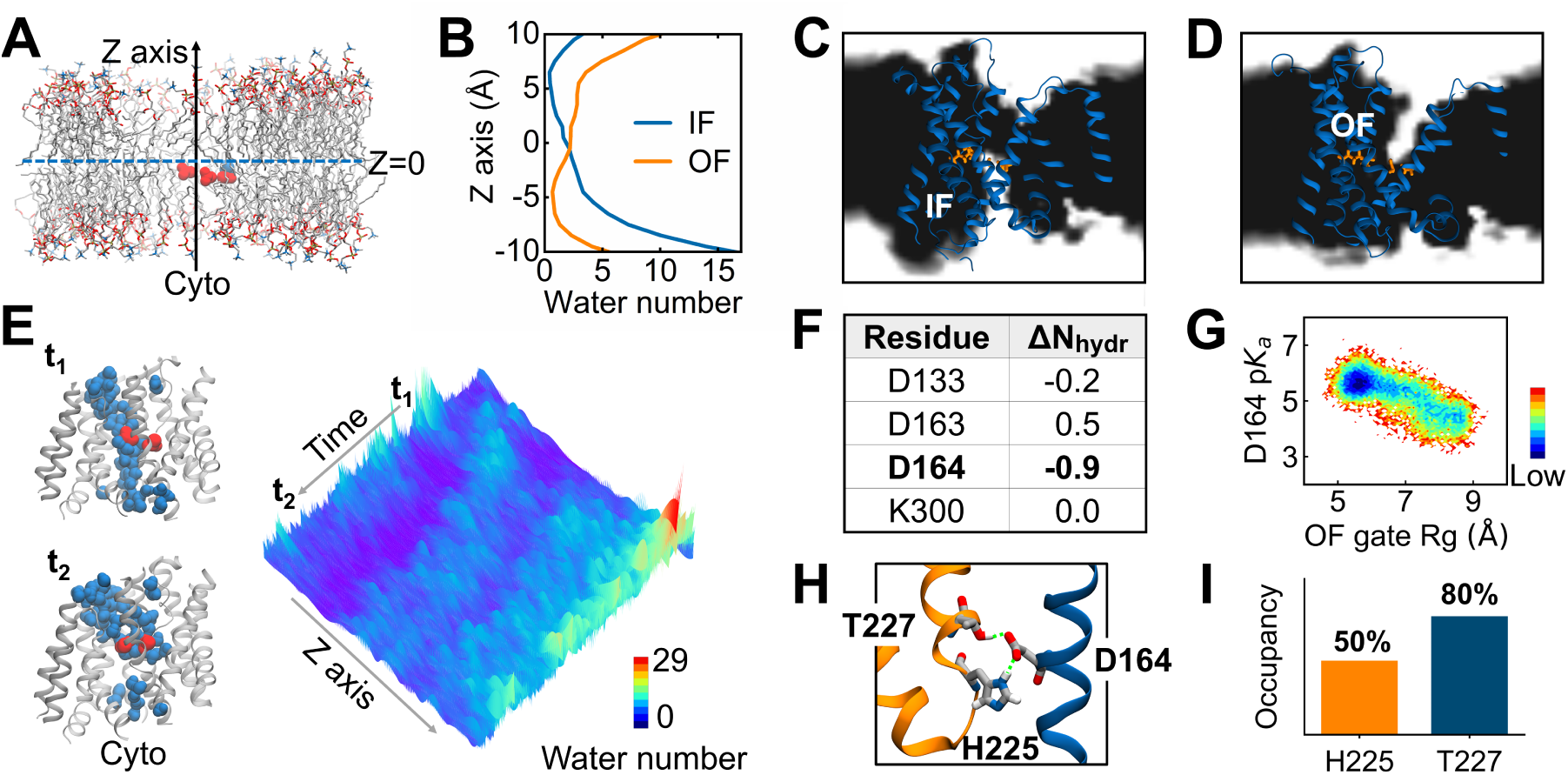
Solvation in response to gate opening. **A** Lipids are displayed with sticks. Side chains of D133, D163, D164 (red) and K300 (blue) are shown with spheres. The solid arrow is the membrane normal along z axis and the dashed arrow perpendicular to the normal is placed at z=0. **B** Water numbers as a function of z coordinate for IF (gray) and OF (orange), respectively. **C** Water density side view of IF. Solvent inaccessible region is colored black. Protein is displayed with the new cartoon model. Orange sticks are the four side chains. **D** Water density side view of OF. **E** Water number as a function of z coordinate and time. Two frames are extracted to show continuous (up) and disrupted (down) water chains (blue spheres), respectively. Protein is presented with the new cartoon model. Residues D163 and D164 are displayed with red spheres. **F** Hydration number changes from IF to OF (ΔN_hydr_) for side chain oxygens of D133/D163/D164 and side chain nitrogen of K300. **G** Free energy map of OF gate Rg (Å) and p*K*_a_ of D164. **H** A snapshot showing the hydrogen bonds (dashed lines) between D164 and H225 and between D164 and T227 in OF. Residues are shown as sticks. **I** Occupancies of the two hydrogen bonds, which are 50% and 80% for D164-H225 and D164-T227, respectively.

Solvation effects on the four ionizable residues of interest were also examined by calculating mean hydration number changes (ΔN_hydr_’s) from IF to OF for Asp133, Asp163, Asp164 and Lys300. Here the hydration number refers to the number of water in the first solvation shell of an ionizable site, where a cutoff distance of 3.5 Å between the water oxygen and side chain nitrogen of Lys or the nearest side chain oxygen of Asp was utilized. As enumerated in Fig. 5F, ΔN_hydr_ decreases by 0.9 for Asp164, indicating that Asp164 in OF is more buried than in IF. In contrast to Asp164, ΔN_hydr_ for Asp163 is 0.5, meaning that Asp163 is more exposed to solvent in OF than in IF. As to Asp133 and Lys300, both are highly buried (Supplementary Table 2) and changes of hydration number are negligible. With nearly one less hydration in OF, p*K*_a_ of Asp164 is supposed to increase. However, from Fig. 5G, p*K*_a_ of Asp164 computed by DeepKa ^44^ decreases from IF to OF, in agreement with that offered by PropKa (Supplementary Fig. 15) ^45^, which implies that the desolvation could be compensated presumably by charge-charge or hydrogen bonding interaction. Indeed, Asp164 in OF forms hydrogen bonds with side chains of His225 and Thr227 in dimer (Fig. 5H), especially the later with a higher occupancy of hydrogen bonding (Fig. 5I), which may contribute to the stabilization of OF structure. Here the cut off distance between the donor hydrogen and the acceptor of a hydrogen bond is 2.4Å ^46^.

### Sodium-coupled Na^+^/H^+^ exchange

A previous computational study has pointed out that sodium would follow the path of water entering the funnel and eventually bind with active site residues, including backbone of Thr132 and side chains of Asp163 and Asp164 ^24^. However, the recognition is restricted by IF and three microseconds at most of MD simulations ^24, 25^. In this work, we made it possible to comprehensively explore the mechanism of sodium binding with both IF and OF accessible. Besides, the simulation length has been extended to ten microseconds.

To explore possible sodium binding modes, probabilities of sodium bound to individual residues were calculated as a function of time. When Lys300 is protonated in IF (+/IF), sodium initially binds to Asp164, then dissociates from this residue, which takes approximately ten microseconds, and simultaneously turns to interact with Thr132 and Asp163 (Fig. 6A). Once Lys300 is neutralized in IF (0/IF) (Fig. 6B), an additional sodium binding site is provided by Lys300, where a sodium ion has been found directly coordinated by the lone pair electrons of Lys300 epsilon-amino nitrogen in a recent MD study in IF by Winkelmann et al ^25^. Likewise, sodium is firstly received by Asp164 and then delivered to Thr132, Asp163 and Lys300 immediately. On the other hand, sodium in OF is primarily coordinated by Thr132, Asp163 and neutralized Lys300 after 2 microseconds (Fig. 6C and D), in line with that in IF. Notably, the equilibration of sodium in the active site is slower for Lys300 charged, which is acceptable as sodium would be expelled by positively charged Lys300. In addition, above data indicate that sodium entering the funnel is spontaneous and fast (Supplementary Table 3), which agrees with previous CpHMD ^24^ and MD ^25^ simulations.

**Figure 6:**
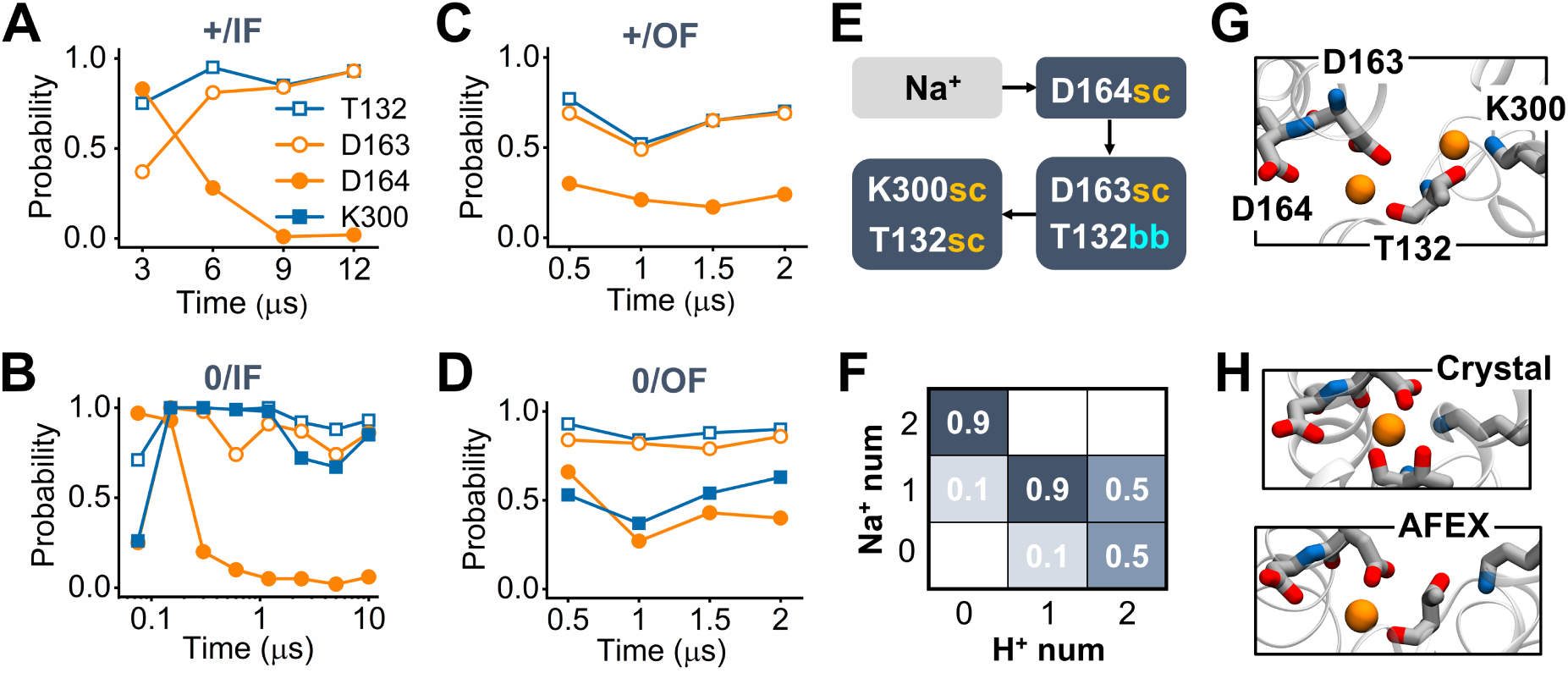
Sodium binding modes. **A** Binding probabilities for T132 (open square), D163 (open circle) and D164 (solid circle) in IF where K300 is charged (+/IF). **B** Binding probabilities for T132, D163, D164 and K300 (solid square) in IF where K300 is neutralized (0/IF). **C** Binding probabilities for T132, D163 and D164 in OF where K300 is charged (+/OF). **D** Binding probabilities for T132, D163, D164 and K300 in OF where K300 is neutralized (0/OF). **E** A schematic diagram of sodium binding pathway. sc and bb are abbreviations of the sidechain and backbone of a residue, respectively. **F** Probability distribution of sodium number (Na^+^ num) in the active site as a function of proton number (H^+^ num) carried by K300 and D164. Sodium is considered bound if the distance to the nearest carboxylate oxygen of D163/D164, the backbone carbonyl or side chain oxhydryl oxygen of T132 or side chain amine nitrogen of neutralized K300 is below 3 Å . The plot is based on the last 1 microsecond of simulations. **G** A snapshot showing two sodium ions (sphere) bound to the active site residues (stick). **H** Two snapshots showing one sodium ion bound to the active site residues in simulations that started from crystal (Crystal) and AFEX-predicted (AFEX) structures, respectively.

As illustrated Fig. 6E, there are 2 to 3 stops for a sodium ion that enters the active site. The sodium ion interacts with Asp164 first that acts as a transient stop. If Lys300 is charged, then the terminal stop is Asp163 side chain together with Thr132 backbone (Supplementary Fig. 16). Otherwise, the sodium ion will be further passed to Lys300 and Thr132 side chains (Supplementary Fig. 16). In that case, sodium binding sites, Asp163 side chain and Thr132 backbone, are vacant. Thus, it is of interest to see whether an additional sodium ion is allowed to bind. As shown in Fig. 6F, when the two putative proton carriers, namely Asp164 and Lys300, are deprotonated, a pair of sodium ions are found in the active site, which was also observed by Winkelmann et al. in their simulations ^25^. Next, Lys300 is protonated and Asp164 remains deprotonated. It is found that only one sodium ion is bound. Surprisingly, when both residues are protonated, there is still one sodium ion bound but with lower occupancy of 50%. In fact, the averaged number of sodium ions in the active site, especially when Asp163 and Lys300 are both protonated, is higher in IF than that in OF (Supplementary Fig. 17), consistent with the physiological transport of sodium from cytoplasm to periplasm. Basically, above statistics works for both IF and OF, supporting the symmetric transport measured by solid support membrane-based electrophysiological experiment^47^.

Finally, as shown in Fig. 6G, the reaction cavity can accomodate two sodium ions when Lys300 is neutralized (Supplementary Movie 2). The sodium ion that enters the cavity first is ligated by the side chain amine nitrogen of Lys300 and side chain oxhydryl oxygen of Thr132. Now that Lys300 is combined with a sodium ion, the negatively charged gap between TMs IVc and XIp nearby would help enhance sodium binding affinity (Supplementary Fig. 18). As to the secondly loaded sodium ion, it binds with side chain carboxyl oxygens of Asp163 and backbone oxygen of Thr132, resembling the binding configuration in Fig. 6H where Lys300 is charged and likely to interact with either Asp163 side chain or the gap, depending on the initial structure, crystalized or AFEX-predicted, of a simulation.

## Discussion

At present, three crystal structures of NhaA are available ^15, 25, 28^. Unfortunately, they all belong to IF state. Thus, it becomes urgent to uncover the OF state that accounts for the second half of the mechanism. Recently, an OF structure was predicted by enhanced sampling ^41^. Nevertheless, the essential dynamic information would be lacked and the interaction between substrates and protein were missed too. Encouragingly, OF structure was accessed in this work by regular MD simulations from which the indispensable sodium binding event can be elucidated. Notably, these simulations were initialized from the structure predicted by AFEX ^42^. Apart from OF, the well-studied IF state was obtained too. As a result, the AFEX-predicted structure could be identified as a transient state between IF and OF.

Looking into the structure predicted by AFEX, we propose that the disrupted hydrophobic interactions between core residues (Ile168 and Ile345) and Phe72 of dimer enable OF gate to open and the access to OF. More importantly, our simulations suggest that Phe72 plays a key role in determining to what extent core moves with respect to dimer at membrane normal. It is worth noting that once OF or IF was accessed, no backward transition was observed. For one thing, one turnover cycle of NhaA takes about 1 millisecond ^12^, two orders of magnitude higher than the present simulation time scale of ten microseconds. For another, hydrophobic forces in a gate could be overestimated by an additive force field ^49^. Thus, simulations extended to one thousand microseconds or more would be necessary for such a conformational transition. Alternatively, polarizable force fields, such as Drude ^49^, would help reduce hydrophobic interactions in a gate and therefore accelerate gate opening.

The hypothesis that Lys300 carries a proton for substrate exchange puts us forth to investigate possible effects of switching the protonation state of this residue. First of all, based on free energy maps, deprotonation of Lys300 lowers the energy barrier between IF and OF, which would accelerate the conformational dynamics. Particularly, only when Lys300 is neutralized, IF or OF gate can be wide open and simultaneously the other is closed, consistent with earlier CpHMD simulations in IF ^24^. Here we suggest that the neutralization of Lys300 is essential for the conceptual alternating gate opening. On the other hand, when Lys300 is charged, the structure is more likely trapped in either IF or OF state. Previous CpHMD or MD simulations under IF revealed that TM V bends when Lys300 is deprotonated, based on which Shen et al. suggested that TM V bending may precede the transition from IF to OF ^24^. Intriguingly, such a conformational change is observed too by current simulations where Lys300 is charged in OF. As a result, our simulations further suggest TM V bending could be the precedence of large conformational transitions between IF and OF.

To explore the mechanism of NhaA, cofactors from solution, such as water molecules and salt ions, should be taken into account. Surprisingly, a continuous water chain, though short-lived, was formed at the beginning of the AFEX-predicted structure converting to OF. Noting that protons could be leaked through a water wire using the Grotthuss mechanism, which is undesired for this protein, we suggest that the water wire is more likely a result of the thermal fluctuation at the early stage of a MD simulation. As water reach the active site, Asp164, the other proton donor, was less hydrated or more buried in OF than IF. Unlike a hydrophobic environment in IF, Asp164 in OF participated in two hydrogen bonds with His225 and Thr227 in dimer, respectively. Accordingly, we hypothesize that OF state could be stabilized by the two hydrogen bonds between core and dimer, breaks of which would facilitate Asp164 to uptake a proton at periplasm and simultaneously the conformational change from OF back to IF.

One advantage of current MD simulations is that substrate ion binding can be depicted explicitly. It has been a consensus that two protons are exchanged for one sodium ion in a transport cycle. Normally one would expect a single sodium ion bound to the active site. However, our simulations show that the active site is capable of accommodating two sodium ions if Lys300 is neutralized, consistent with a recent MD study of NhaA in IF by Winkelmann and coworkers ^25^. However, in our simulations, two sodium ions were harbored by Thr132, Asp163 and neutralized Lys300, in contradiction with the binding mode found by Winkelmann et al. where Asp164 is one of the binding sites. Based on the present ten-microsecond-timescale simulations in IF, we suggest the binding mode observed by Winkelmann et al. should correspond to the initial stage of the second sodium ion bound to the active site. When Lys300 is charged, one sodium ion is excluded and the other that originally binds with Lys300 and Thr132 sidechains is switched to Asp163 side chain and Thr132 backbone.

It has been reported that positively charged Lys300 on TM X can be attracted by negatively charged Asp163 on TM V via salt bridging^25, 28^ or the cleft capped by terminals of TMs IVp and XIc through hydrogen bonding ^15^, which is essential for the structural stabilization of core. Thus, neutralizing Lys300 would reduce TM X-involved helix-helix interactions and the resulting destabilization of structure, which has been demonstrated by mutating Lys300 to Ala or Cys that has a neutral and short side chain ^31^. Nevertheless, when Lys300 is neutralized, the structure remains stable in simulations, which could be explained by the positive charge offered by a sodium ion bound to the side chain nitrogen of Lys300.

Here we propose that the additional sodium ion may act as a counter ion that stabilizes the negatively charged gap between TMs IVp and XIc in the active site when Lys300 is neutralized. It is worth noting that bile acid transporters with a LeuT-like fold also accommodate two sodium ions. Specifically both sodium ions are embedded separately in the gaps formed by two discontinuous helices of a LeuT transporter ^50^, which is analogous to the clefts created by crossed TMs IV and XI of NhaA and occupied by Asp133 and Lys300, respectively ^2, 15^. Furthermore, it is found that one of the two sodium ions is substituted by the *ɛ*-amino group of a lysine side chain in ApcT, a homology of LeuT ^51^, which may help validate the present Lys300-carried proton replaceable by a sodium ion to maintain local charge balance. Back to NhaA, it should be emphasized that two sodium ions are bound to the side chain and backbone of Thr132, respectively, implying that Thr132 might play a central role in sodium binding. Finally, given protonated Lys300, further neutralization of Asp164 diminishes but does not inhibit sodium binding to Asp163 and Thr132, especially in IF, which to some extent highlights the importance of a negative charge on Asp163 and therefore supports the observation that neutralizing mutation of Asp163 abolishes sodium binding ^33^.

In summary, the present work aims to reveal the transport mechanism of NhaA under the hypothesis that Lys300 is one of the proton carriers. First of all, as illustrated in Fig. 7A, the protein in view of energy was placed by AFEX to a barrier between IF and OF, which allows the access to either IF or OF. Further deprotonation of Lys300 switches the bias of accessing from OF to IF, though the biological relevance is not clear, and simultaneously smooths the barrier, which would accelerate the kinetics. Next, as elucidated in Fig. 7B, we propose the following gating mechanism of NhaA where Phe72, a OF gate residue in dimer, plays a key role. When NhaA is in resting state, both IF and OF gates are closed. Once activated, IF and OF gates open alternately. Notably, OF gate opening is accompanied by a large conformational change of core with respect to dimer that includes an angular and simultaneously an elevator-like movements restrained by Phe72. Finally, we come up with a novel mechanism for sodium-proton antiporting. As shown in Fig. 7C, putative proton carriers Asp164 and Lys300 are ionizable while Asp163 fixed in charged state. Initially Adp164 and Lys300 are protonated in resting state. Asp163 forms a salt bridge with Lys300. When activated, distinct sodium binding modes for IF and OF are applied. In IF, Asp164 and Lys300 release their protons to cytoplasm. Two sodium ions from cytoplasm enter the active site successively. Particularly, one sodium ion is coordinated by Asp163 side chain and Thr132 backbone and the other is bound to side chains of Lys300 and Thr132. Switching to OF, Asp164 and Lys300 are protonated. The sodium ion that combines with Asp163 and Thr132 is excluded and released to periplasm. Its place is occupied by the other previously bound with Lys300 and Thr132. Apparently, such a sodium-assisted mechanism preserves electrogenic transport or more specifically the sodium-proton exchange stoichiometry of 1:2 ^10^. In the end, we propose that breaks of two Asp164-involved hydrogen bonds between core and dimer might be essential for Asp164 to uptake a proton from periplasm and the conformational change from OF back to IF.

**Figure 7:**
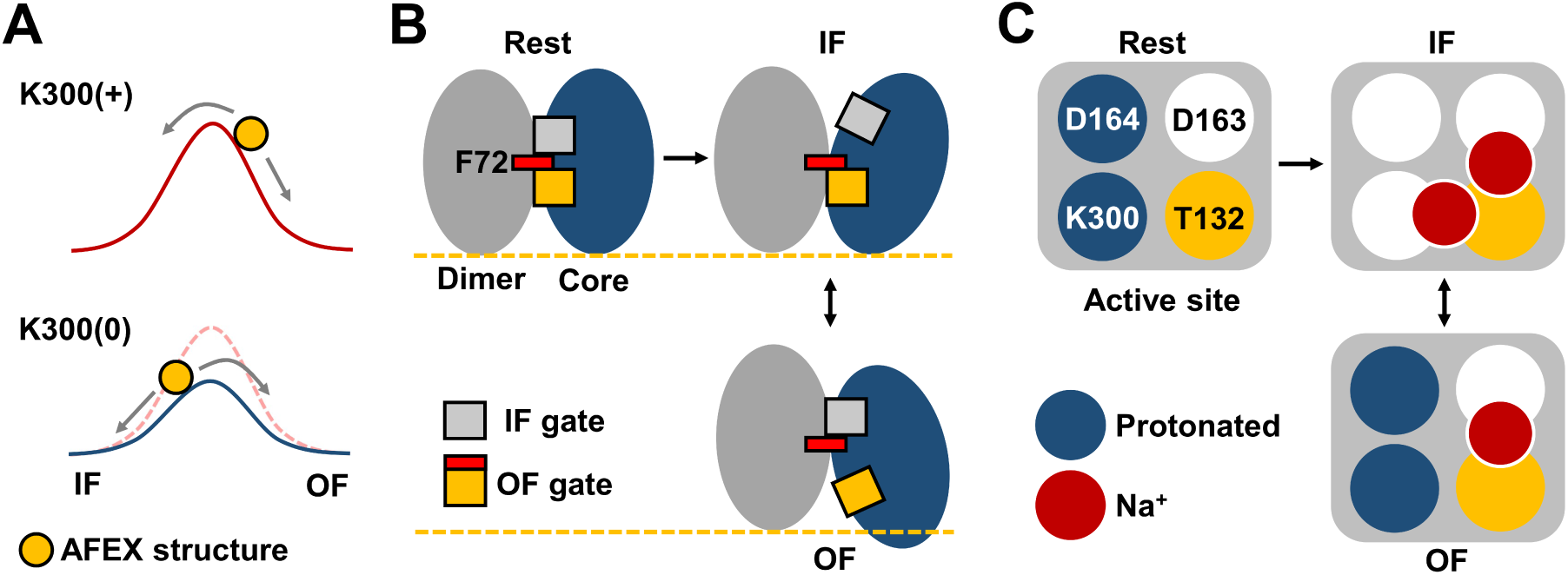
Schematic diagrams of transport mechanism. **A** Energy curves of simulations that started from the AFEX-predicted structure (orange ball) for K300 in charged (K300(+), red) and neutralized (K300(0), blue) states, respectively. Gray arrows indicate the access to IF or OF state. The dashed red curve is a duplication of the solid one above for K300(+). **B** Conformational change of NhaA. The crystal structure measured at acid pH is the inactive state of NhaA (Rest). The arrow from Rest to IF indicates that NhaA is activated at alkaline pH, which is characterized by the transition (double-head arrow) between IF and OF states. Gray and blue ellipses represent dimer and core domains, respectively. gray square is the IF gate. F72 of OF gate is displayed separately by a red rectangle and the rest by an orange square. A dashed line is applied to probe the displacement of the core with respect to the dimer along the membrane normal. **C** A schematic diagram of sodium-assisted sodium-proton exchange. Rectangles filled with gray represent the active site in the core domain. White circles represent deprotonated D163, D164 or K300. T132 is displayed with a orange circle. Circles colored blue indicate that residues are protonated. Red circles represent sodium ions. Overlaps of circles indicate sodium binding.

## Methods

### Alternative structure prediction

AFEXplorer (AFEX) is a generic method that tailors the protein structure prediction model AlphaFold ^48^ to infer alternative conformations by optimizing the input embeddings ^42^. AFEX-predicted structures should satisfy a prior knowledge which is expressed as constraints in a coarse coordinate space. The concept of the constraints is similar to that of collective variable (CV) in the realm of MD simulation, which refers to the generic distance between the generated structures and the prior knowledge. Using the inference and back propagation operation of AlphaFold framework for prediction and gradient-based optimization, respectively, alternative structures that satisfy the prior knowledge can be generated by AFEX. In principle, AFEX is general and applicable to any protein that has a prior knowledge available. For instance, alternative structures of four membrane transporters have been predicted successfully by AFEX where the prior knowledge is the C*α* distance of a residue pair involving in the transition between IF and OF states ^42^. Starting from the AFEX-predicted alternative states, MD simulations may sample more extensive conformational space.

In this work, the OF structure of NapA (PDB code: 4BWZ) ^37^, a homolog of NhaA, was utilized as the prior knowledge. As a result, C*α* distance between Arg81 in TM II and Leu143 in TM IV (*d*) was selected as the CV of NhaA, where a distance of less than *z* = 10 Å is expected for the OF state. The loss function is expressed as 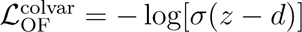, where *σ* refers to the sigmoid function *σ*(*x*) = 1*/*(1 + exp (−*x*)). Finally, the confidence score of predicted structures was normalized and added to the optimized objective function to ensure the quality of predicted structures. After a 500-step optimization, AFEX totally generated 500 predicted structures of NhaA where *d* changes from 26.5 Å to 4.2 Å, along which the states of predicted structures can change gradually. Among these structures with reduced *d*, the one (AFEX.pdb in SI) that owns crystal structure-like closed IF and OF funnels was selected, such that details of a AFEX-predicted structure transiting to the OF state could be captured.

### System preparation

Simulations in this work were prepared by the Membrane Builder module of CHARMM-GUI interface ^52–54^ where the starting structures came from the crystal structure (PDB code: 4AU5) ^28^ and the AFEX-predicted model ^42^. Truncated N- and C-terminals were acetylated and amidated with CH_3_CO and NH_2_, respectively. Protonation states of Asp, Glu, His, and Lys residues under the optimal pH of 8.5 for NhaA were determined based on p*K*_a_’s calculated in our previous work ^24^. It’s found that only Lys300 upon sodium binding is able to sample both protonated and deprotonated states at pH=8.5. As a result, two simulation sets were built up where default protonation states Asp(-)/Glu(-)/His(0)/Lys(+)/Cys(0)/Arg(+) were applied to the first set. Whereas in the second set Lys300 was fixed in its neutral state Lys(0). Eight extra simulations that started from equilibrated IF and OF were set up, in which both Asp164 and Lys300 were protonated to mimic NhaA uptaking two protons from periplasm and the status before sodium-proton exchanging at cytoplasmic side. The default rectangular box type was selected and an initial guess of box length at x or y axis was set 80 Å . The protein was inserted in a lipid bilayer composed by 1-palmitoyl-2-oleoyl-*sn*-glycero-3-phosphocholine (POPC) lipids. Then box length at z axis was determined according to the default water thickness of 22.5 Å capped to both sides of the protein in z direction. Besides, water molecules were added to fill the pore if any in the protein. Finally, sodium and chloride ions were added to neutralize the simulation system and reach the physiological ionic strength of 0.15 M.

### MD simulations

CHARMM36m ^55^ and CHARMM36 ^56^ force fields were utilized to describe the protein and lipids, respectively. The atom-pair-specific adjustment NBFIX was applied to sodium and chloride ions, which would attenuate the overestimated binding affinity of sodium to carboxylates ^57^. CHARMM-modified TIP3P model was used to represent water ^58, 59^. Simulation systems were relaxed or equilibrated following the multi-step protocol in CHARMM-GUI. Then simulations that started from the AFEX-predicted structure lasted 500 nanoseconds, except for the four extended to 2 microseconds. On the other hand, two simulations initialized from the crystal structure lasted 12 and 10 microseconds for protonated and deprotonated Lys300, respectively. The eight additional simulations each lasted 2 microseconds. As a result, the accumulative running length reaches 64 microseconds. All simulations were performed in the NPT ensemble using the OpenMM package ^60^. The Hoover thermostat ^61^ was employed to keep the temperature of 310 Kelvin. The Langevin piston coupling method ^62^ was applied for the constant pressure of 1 atm. Under the periodic boundary condition, the particle mesh Ewald method was used to compute electrostatic interaction where the real-space cutoff is set 12 Å ^63^. As to the reciprocal space, the 1-Å grid spacing and sixth-order spline interpolation were applied. The force switch scheme was employed to deal with the van der Waals interaction where the switching on and off distances are 10 and 12 Å, respectively. The use of SHAKE algorithm ^64^ allows a time step of 2 fs.

### Analysis

Unless otherwise stated, the first 100 nanoseconds of the present simulations were discarded for final trajectory analyses by VMD ^65^ or MDAnalysis ^66^. p*K*_a_’s were estimated by two distinct methods, namely the deep learning-baed DeepKa ^44^ and an empirical formula PropKa 3.0^45^. We note that PDB is the only input format readable by DeepKa or PropKa. Frames of a simulation were saved by VMD as PDB format files for p*K*_a_ calculations. Finally, structures were visualized by VMD.

## Data availability

The data that support this study are available from the corresponding authors upon request. All files required for producing the trajectories of the MD simulations can be found at repository: https://gitlab.com/computational-biophysics/nhaa/input.

## Acknowledgements

Y.H. acknowledges the support of the National Natural Science Foundation of China (11804114) and the Natural Science Foundation of Fujian Province, China (2023J01329). Financial support is provided by the National Natural Science Foundation of China (32171247) and the Pioneer and Leading Goose R&D Program of Zhejiang (2023C03109, 2024SSYS0036) to J.Huang.

## Author contributions

These authors T.X. and J.He contributed equally to this work. Y.H. and J.Huang designed the project. X.T. and J.He performed MD simulations. X.T. and J.Huang provided the initial structure predicted by AFEX. Y.H., J.He, S.S., T.X. and Y.C. conducted data analysis. Y.H. wrote the first draft and J.Huang helped finalize the manuscript. All authors commented on the manuscripts.

## Competing Interests

The authors declare no competing financial interests.

## Additional information

### Supplementary material

Supplementary Information accompanies this paper.

## Notes

### Competing Interest Statement

The authors have declared no competing interest.

