## Supporting Information for "Molecular mechanism of Na^+^/H^+^ antiporting in NhaA"

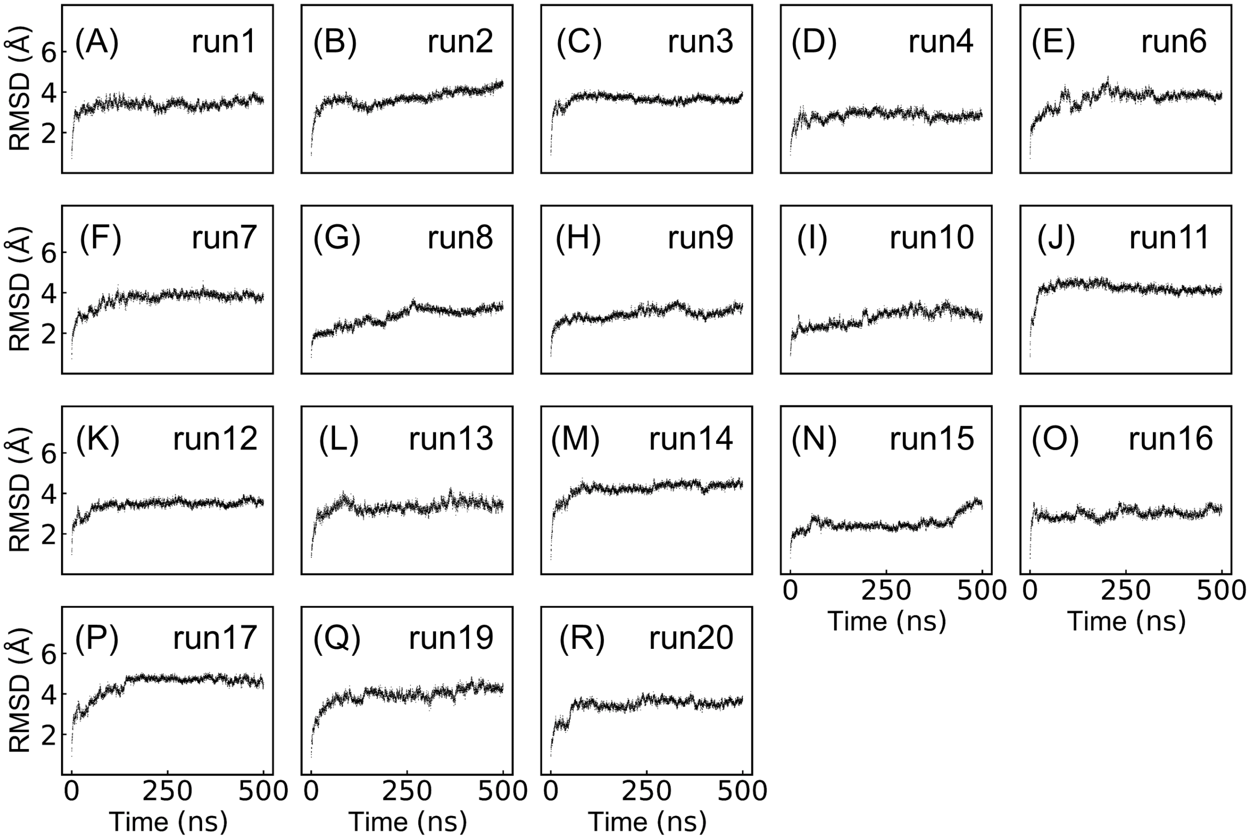


Figure 1: (A)-(R) Root mean square deviation (RMSD) of 18 simulations that started from the AFEX-predicted structure. Here K300 is charged or K300(+).

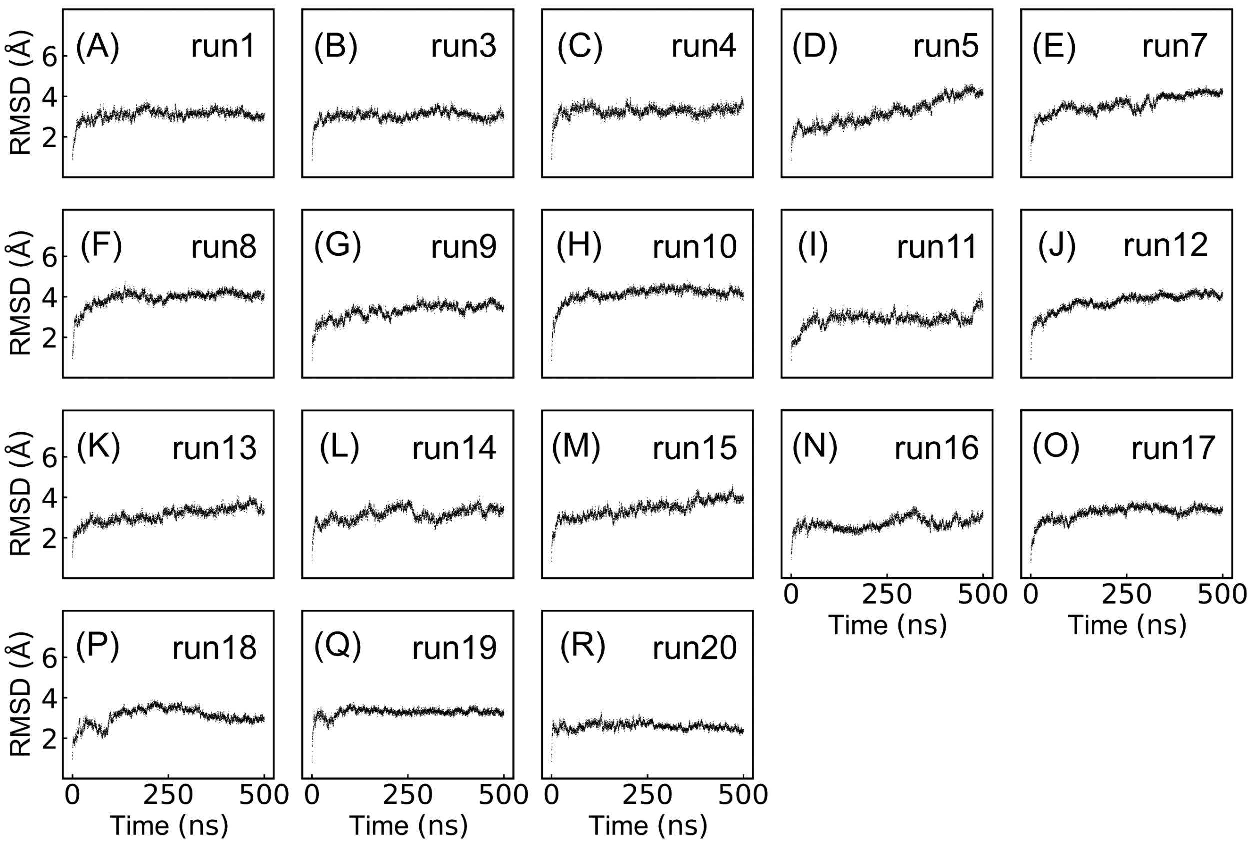


Figure 2: (A)-(R) RMSD of 18 simulations that started from the AFEX-predicted structure. Here K300 is neutralized or K300(0).

Table 1: Minimal distances (Å) between OF gate side chains in the crystal (PDB code: 4AU5) and AFEX-predicted structures.

| Residue pair | $d_{4AU5}$ | $d_{AFEX}$ | $\Delta$ |
|---|---|---|---|
| F72-A167 | 8.1 | 7.9 | -0.2 |
| F72-I168 | 3.7 | 5.2 | 1.5 |
| F72-F344 | 5.3 | 5.2 | -0.1 |
| F72-I345 | 6.8 | 8.2 | 1.4 |
| I168-I345 | 3.56 | 8.26 | 4.7 |

[a] $\Delta = d_{AFEX} - d_{4AU5}$

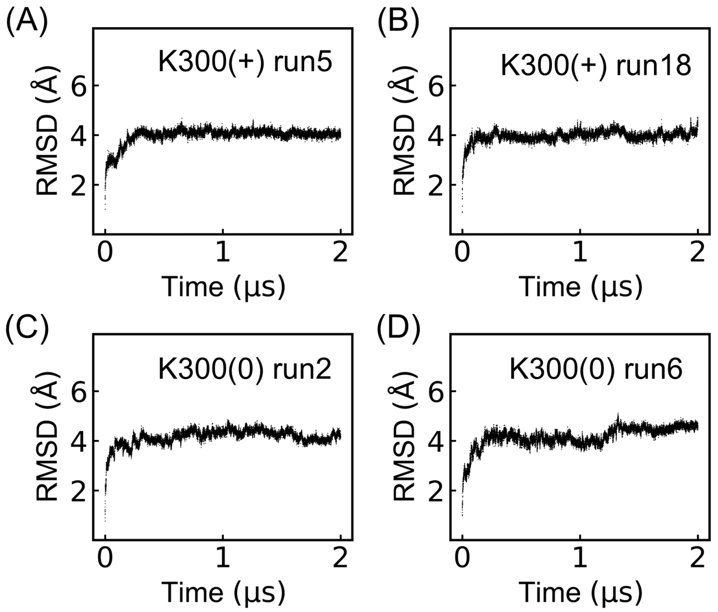


Figure 3: (A)-(B) RMSD of simulations K300(+)run5 and K300(+)run18. (C)-(D) RMSD of simulations K300(0)run2 and K300(0)run6. Above simulations started from the AFEX-predicted structure and relaxed to OF state.

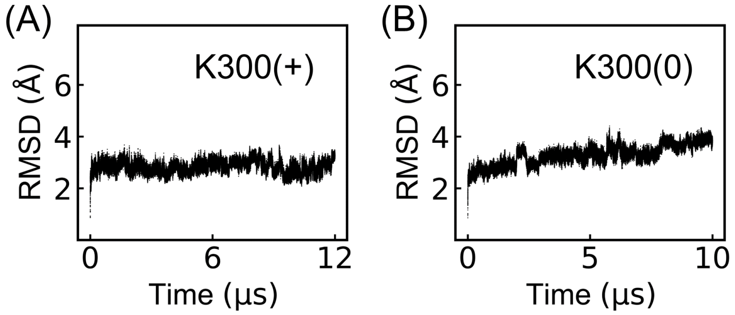


Figure 4: RMSD of simulations that started from the crystal structure (PDB code: 4AU5). (A) K300(+). (B) K300(0).

Table 2: Hydration number of carboxyl oxygens on D133/D163/D164 or sidechain nitrogen on K300 in IF/OF state

| Residue | $N_{hydr}$(IF) | $N_{hydr}$(OF) | $\Delta N_{hydr}$[a] |
|---|---|---|---|
| D133 | 2.8 | 2.6 | -0.2 |
| D163 | 4.4 | 4.9 | 0.5 |
| D164 | 5.5 | 4.6 | -0.9 |
| K300 | 0.8 | 0.8 | 0.0 |

[a] $\Delta N_{hydr}=N_{hydr}$(OF)-$N_{hydr}$(IF)

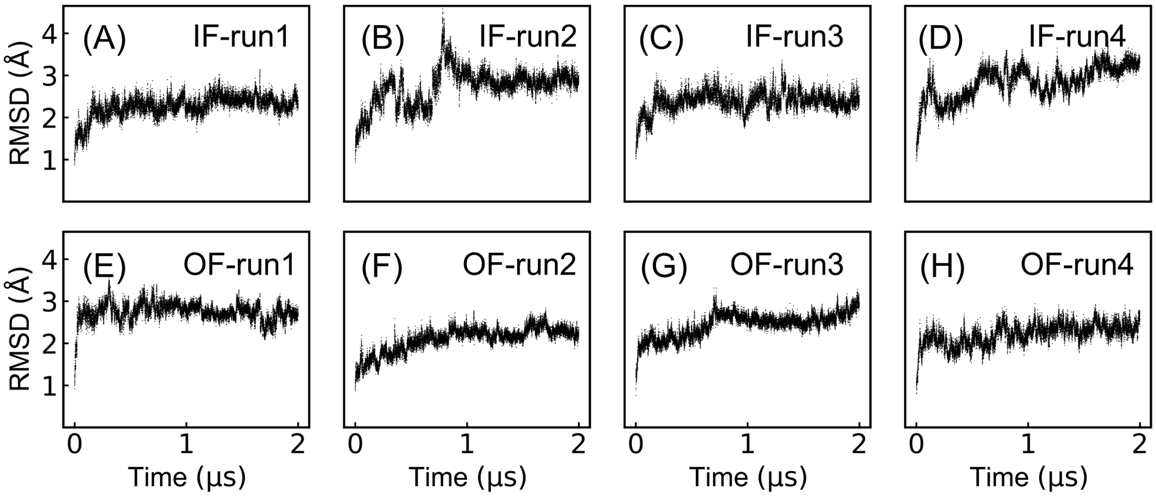


Figure 5: (A)-(H) RMSD of simulations that started from simulated IF and OF structures. K300 and D164 are both protonated.

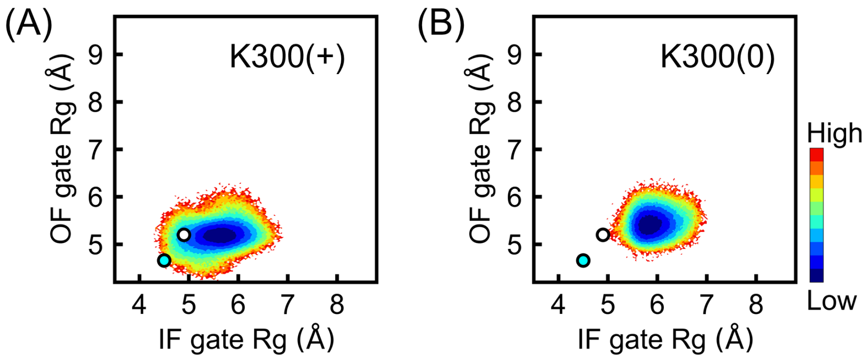


Figure 6: Energy maps of IF and OF gate Rg by simulations that started from the crystal structure (PDB code: 4AU5). (A) K300 is charged or K300(+). (B) K300 is neutralized or K300(0). Positions of the crystal and AFEX-predicted structures are marked by cyan and white circles, respectively.

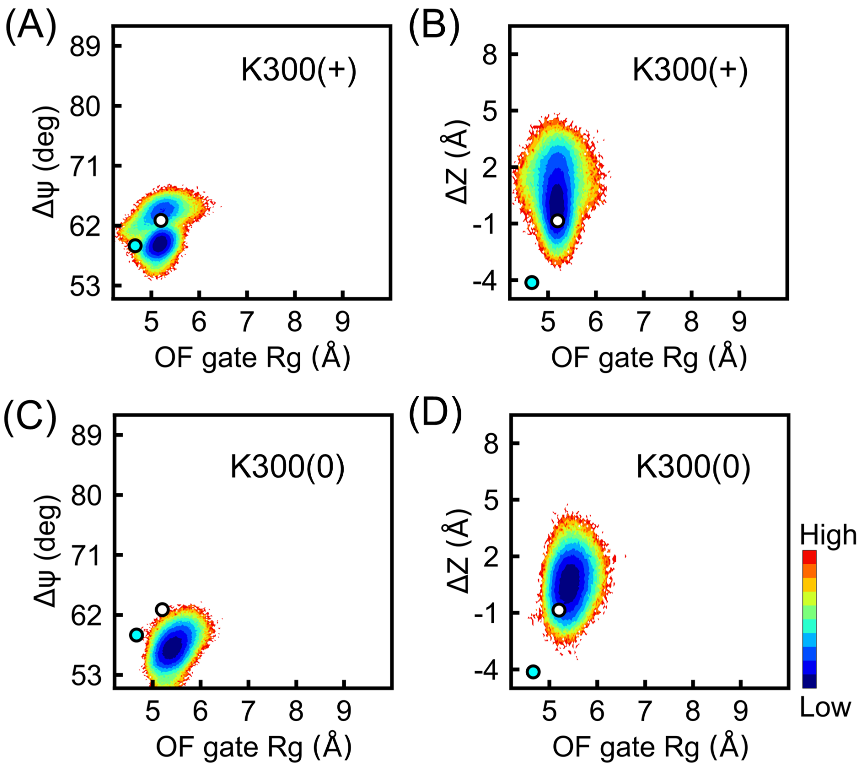


Figure 7: Energy maps of OF gate Rg and $\Delta\Psi$ (or $\Delta Z$) by simulations that started from the crystal structure (PDB code: 4AU5). (A)-(B) K300(+). (C)-(D) K300(0). Positions of the crystal and AFEX-predicted structures are marked by cyan and white circles, respectively.

Table 3: The time taken by the first sodium ion bound to D164 in the forty independent runs that started from the AFEX-predicted structure

| Run number | Time (ns) | |
|---|---|---|
| | K300(+) | K300(0) |
| 1 | 24.2 | 5.8 |
| 2 | 7.9 | 27 |
| 3 | 11.2 | 17.7 |
| 4 | 12.4 | 9.1 |
| 5 | 14.9 | 4.7 |
| 6 | 46.4 | 7.6 |
| 7 | 11.8 | 9 |
| 8 | 288.9 | 95.2 |
| 9 | 165.7 | 12.9 |
| 10 | 20.6 | 10.1 |
| 11 | 31 | 1.1 |
| 12 | 17.7 | 7.2 |
| 13 | 19.3 | 3.5 |
| 14 | 77.5 | 4.4 |
| 15 | 103.6 | 1.3 |
| 16 | 13.7 | 8 |
| 17 | 32.2 | 11.1 |
| 18 | 17.3 | 24.8 |
| 19 | 24.4 | 11.3 |
| 20 | 54.1 | 11 |
| Mean | 49.7 | 14.1 |
| Standard deviation | 68.3 | 20.2 |
| Minimum | 7.9 | 1.1 |
| Maximum | 288.9 | 95.2 |

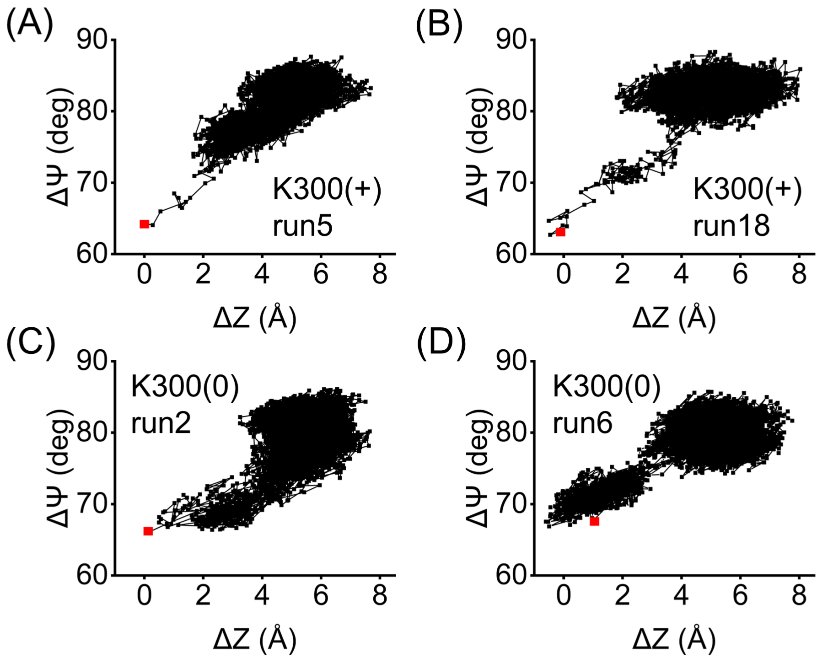


Figure 8: (A)-(B) Trajectories in the correlation plots of $\Delta\Psi$ and $\Delta Z$ for run5 and run18 that relaxed to OF under K300(+). (C)-(D) Trajectories in the correlation plots of $\Delta\Psi$ and $\Delta Z$ for run2 and run6 that relaxed to OF under K300(0). Simulations above started from the AFEX-predicted structure and lasted 2 microseconds. Initial frames are indicated by red squares.

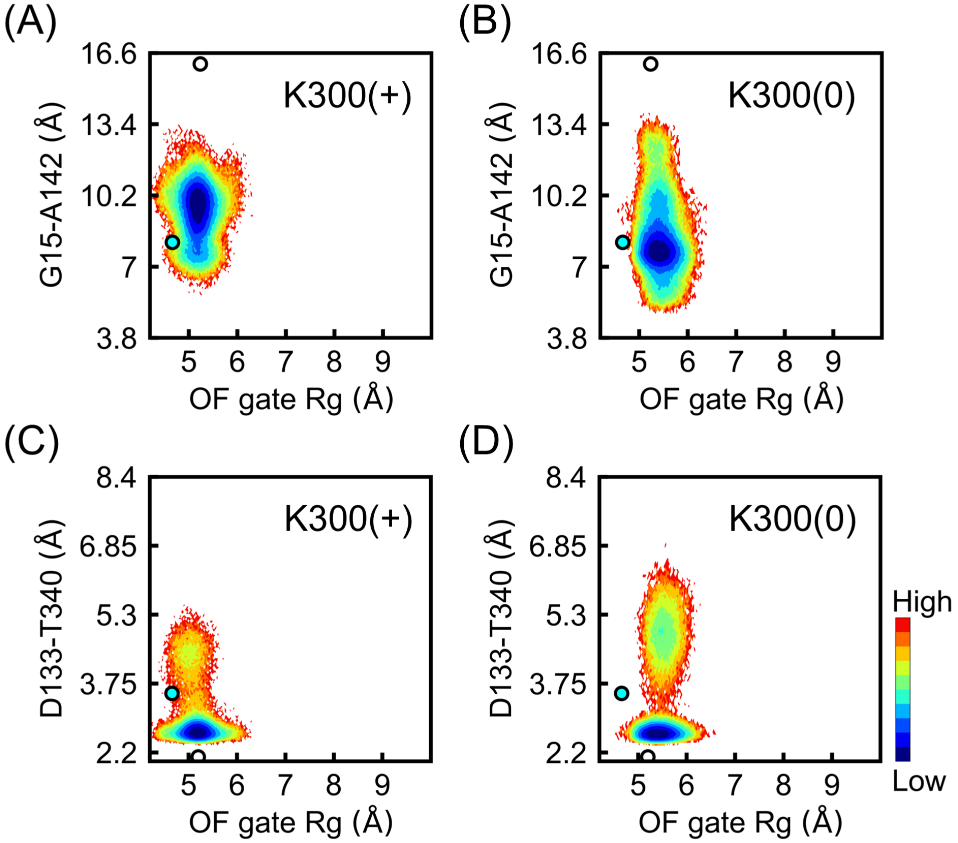


Figure 9: (A)-(B) Energy maps of OF gate Rg and the minimal distance between G15 and A142 (G15-A142) for K300(+) and K300(0), respectively. (C)-(D) Energy maps of OF gate Rg and the minimal distance between D133 and T340 (D133-T340) for K300(+) and K300(0), respectively. Above maps are based on simulations that started from the crystal structure (PDB code: 4AU5). Positions of the crystal and AFEX-predicted structures are marked by cyan and white circles, respectively.

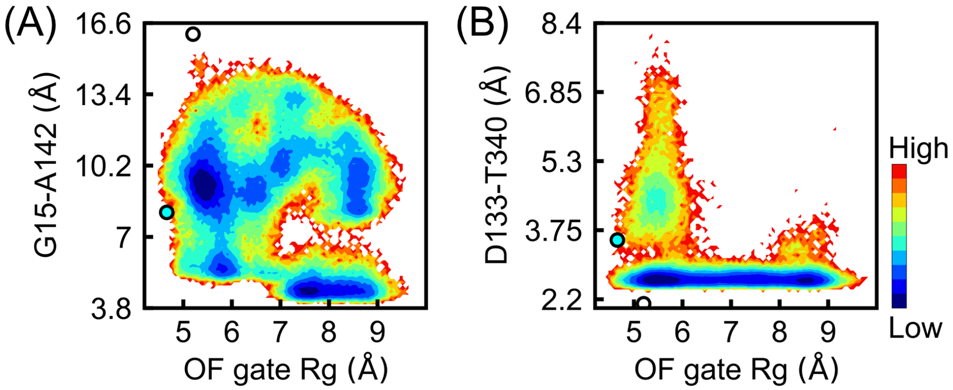


Figure 10: (A) The energy map of OF gate Rg and the minimal distance between G15 and A142 (G15-A142). (B) The energy map of OF gate Rg and the minimal distance between D133 and T340 (D133-T340). Above maps are based on simulations that started from the AFEX-predicted structure. Positions of the crystal and AFEX-predicted structures are marked by cyan and white circles, respectively.

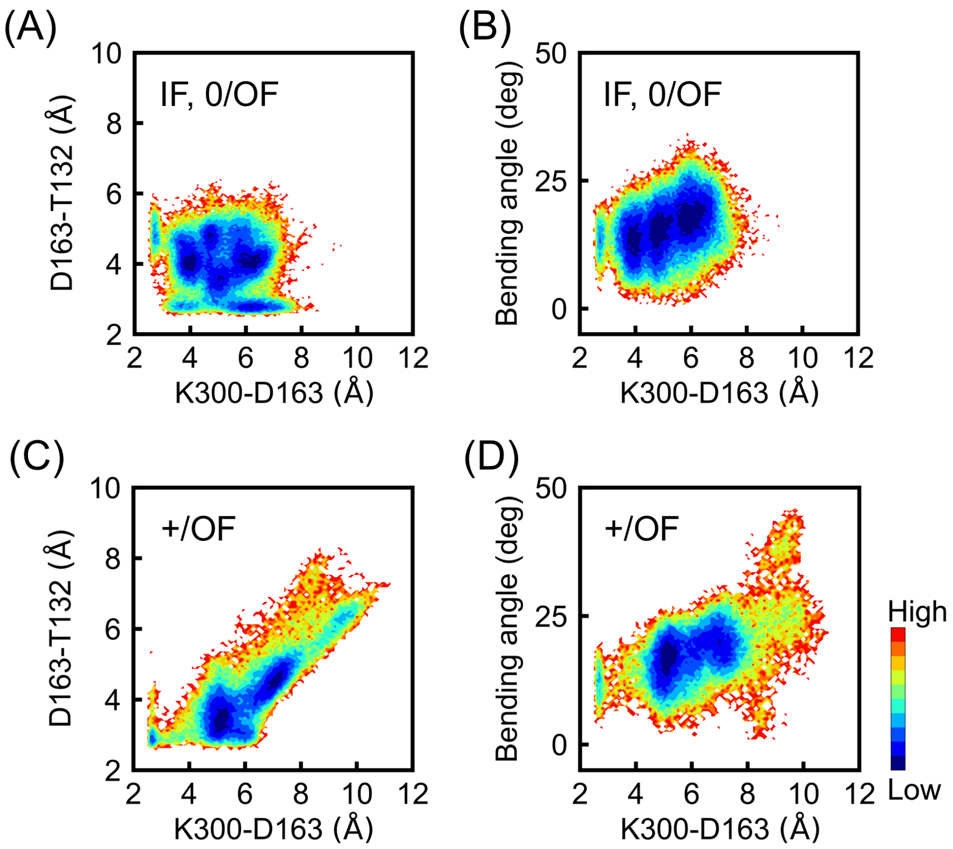


Figure 11: (A) and (C) Energy maps of the minimal distance between K300 side chain nitrogen and D163 side chain carboxyl oxygens and that between D163 side chain carboxyl oxygens and T132 backbone oxygen. (B) and (D) Energy maps of the minimal distance between K300 side chain nitrogen and D163 side chain carboxyl oxygens and TM V bending angle with the unit of degree. The bending angle is defined as the deviation from linearity of the angle formed by the C$\alpha$ atoms of L152, D163 and Phe174 on TM V. (A) and (B) are based on frames in OF where K300 is neutralized and IF. (C) and (D) are based on frames in OF where K300 is charged. Frames are from the 40 independent runs that started from the AFEX-predicted structure.

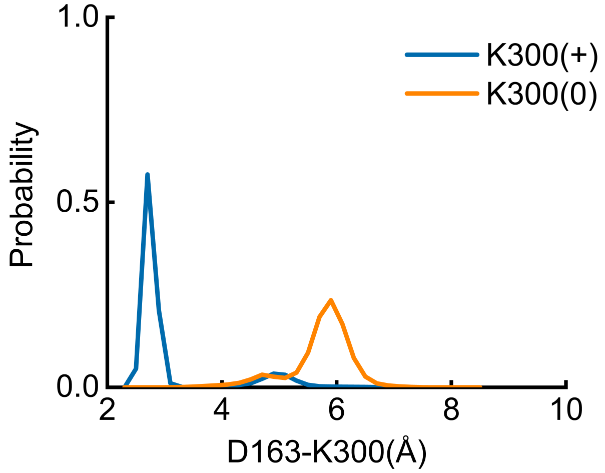


Figure 12: Probability distributions of the minimal distance between D163 side chain oxygens and K300 side chain nitrogen under K300(+) (blue) and K300(0) (orange), respectively. Above analyses are based on simulations that started from the crystal structure (PDB code: 4AU5).

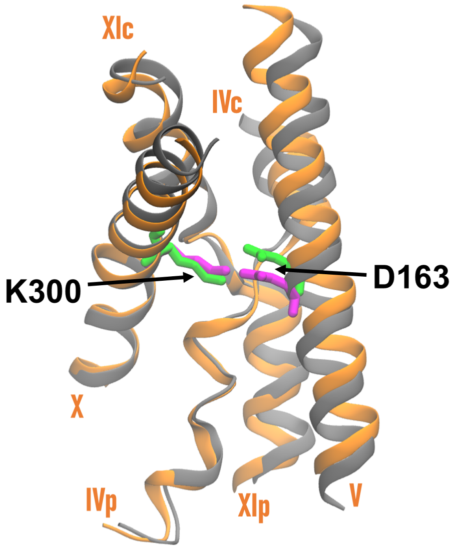


Figure 13: The core domain of the AFEX-predicted structure (orange) is aligned to that of the crystal structure (PDB code: 4AU5, gray). TMs IVc, IVp, V, X, XIc and XIp are shown with the new cartoon model. K300 and D163 for the predicted (green) and crystal (magenta) structures are displayed with sticks.

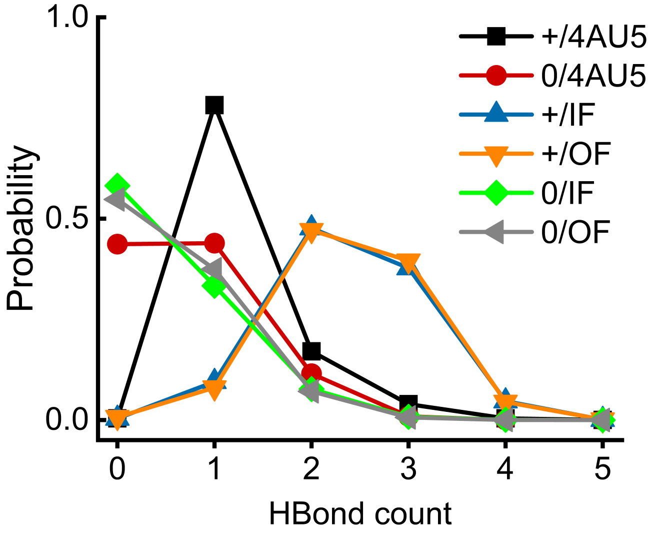


Figure 14: Probability distributions of the hydrogen bond number between K300 sidechain and the cluster composed by backbones of P129, A130, A131, G332 and C335. (+/4AU5) and (0/4AU5) indicate simulations that started from the crystal structure (PDB code: 4AU5) where K300 is charged (black) and neutralized (red), respectively. (+/IF) and (0/IF) are frames in IF where K300 is charged (blue) and neutralized (green), respectively. (+/OF) and (0/OF) are frames in OF where K300 is charged (orange) and neutralized (gray), respectively. Frames applied to the analyses of +/IF, 0/IF, +/OF and 0/OF are extracted from simulations that started from the AFEX-predicted structure.

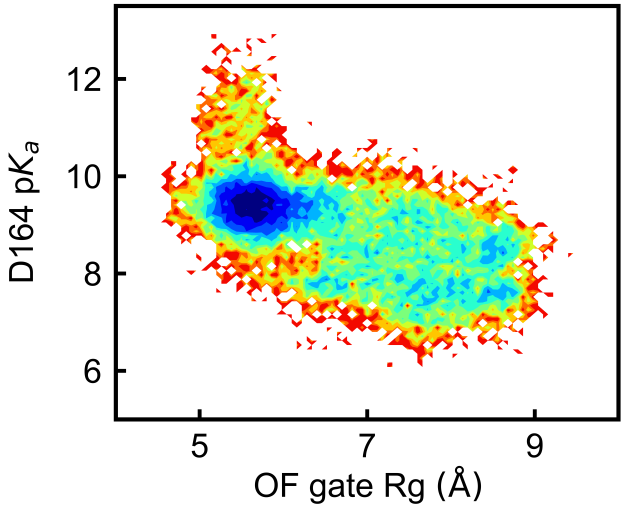


Figure 15: Free energy map of OF gate Rg (Å) and p$K_\mathrm{a}$ of D164 by PropKa.

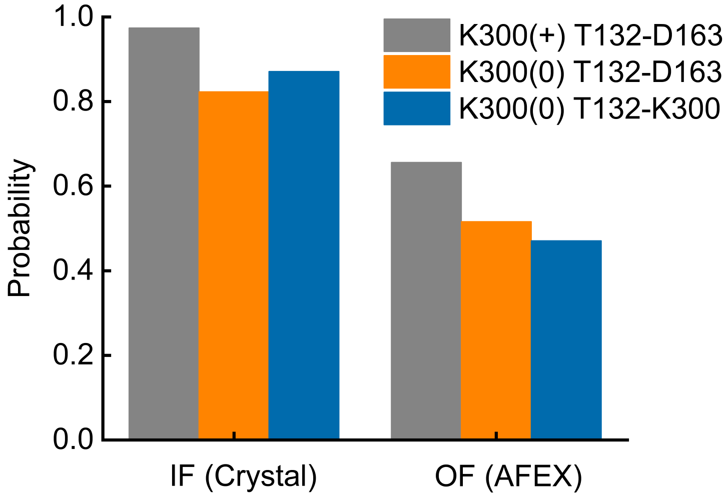


Figure 16: Probabilities of two sodium binding modes in IF and OF states. A sodium ion bound with T132 backbone and D163 sidechain is the first binding mode (T132-D163), where gray and orange histograms correspond to K300(+) and K300(0), respectively. As to the second mode (blue), a sodium ion is coordinated with sidechains of T132 and K300 (T132-K300). Analyses for IF are based on the last 1 microsecond of simulations that started from the crystal structure (PDB code: 4AU5). Analyses for OF are based on the second microsecond of the four runs initialized from the AFEX-predicted structure and extended to 2 microseconds. Sodium is considered bound if the minimal distance from D163 side chain, T132 and K300 side chain is below 3 Å.

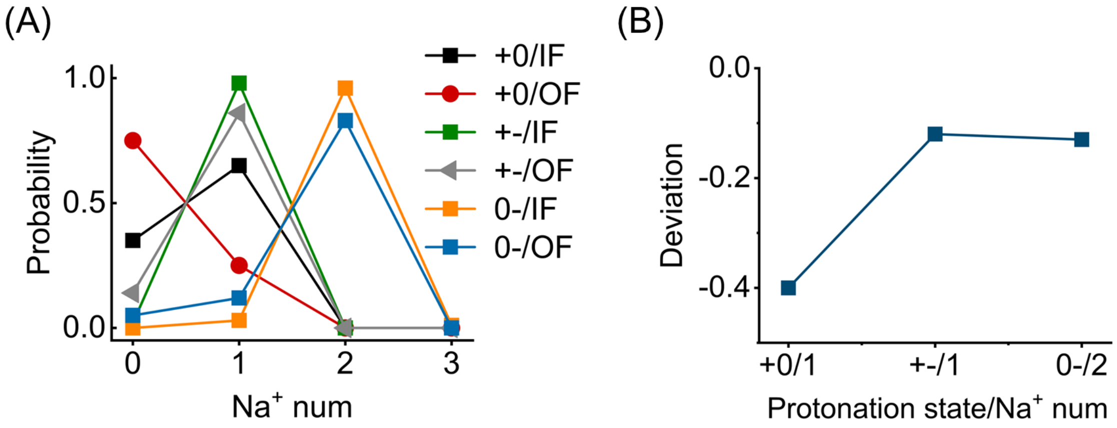


Figure 17: (A) Probability distributions of the number of sodium ions in the active site ($Na^+$ num). There are three combinations of protonation states for K300 and D164, namely +0, +-, and 0-. The first symbol, plus or zero, indicates positively charged or neutralized K300. The second symbol, zero or minus, indicates neutralized or negatively charged D164. States IF and OF are indicated behind the symbols. (B) Deviations of $Na^+$ num in OF from that in IF for +0, +- and 0-, respectively. +0/IF and +0/OF are based on the 8 additional simulations that started from the AFEX-predicted structure. +-/IF and 0-/IF are from the two simulations initialized from the crystal structure. +-/OF and 0-/OF are from four selected simulations that started from the predicted structure and converged to OF state. Sodium is considered bound if the minimal distance from D163 side chain, T132 and K300 side chain is below 3 Å.

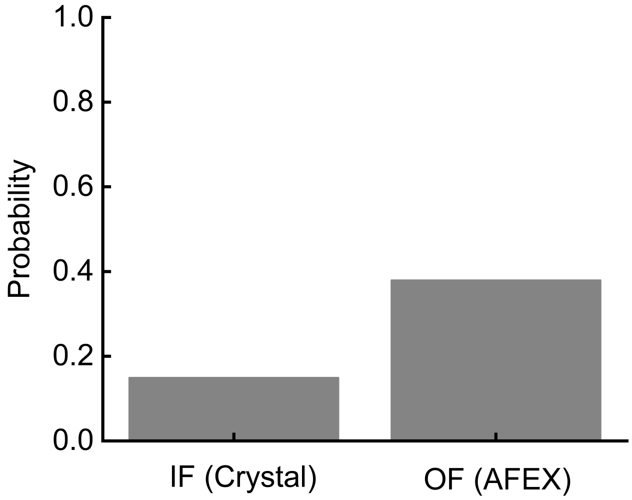


Figure 18: Probabilities of sodium bound with backbones of P129, A130, A131, G332 and C335 that form the negatively charged cleft between TMs IVp and XIc. The analysis for IF are from the last 1 microsecond of simulations that started from the crystal structure (PDB code: 4AU5). The analysis for OF are based on the second microsecond of the four runs initialized from the AFEX-predicted structure and extended to 2 microseconds. Sodium is considered bound if the minimal distance from these backbone oxygens is below 3 Å.
